# Gating Mechanism of the Human Connexin 45 Gap Junction Channel

**DOI:** 10.64898/2026.05.04.722611

**Authors:** Shiv K. Sah-Teli, Evans Eshriew, Amiel Abettan, Ashwini Kedari, Esa-Pekka Kumpula, Jeffrey Saffitz, Vivek Sharma, Juha T. Huiskonen

## Abstract

Gap junction channels formed by the 21-member human connexin family enable direct intercellular exchange of ions and small signaling metabolites, coordinating electrical coupling across cardiac, neural and epithelial tissues. Connexin 45 (Cx45), encoded by GJC1, mediates impulse conduction in the atrioventricular node, His bundle, and Purkinje fibers, where disease-linked mutations cause progressive atrioventricular block and familial atrial fibrillation, yet no experimental structure has been reported, and its regulatory mechanism remains undefined. Here, we determine the structural basis of Cx45 gating and Ca^2+^ regulation using cryo-electron microscopy, mutational analysis, and molecular dynamics simulations. Cryo-EM structures of the apo (2.76 Å), Ca^2+^-bound (2.65 Å), and E41A mutant (3.55 Å) channel reveal a neck constriction formed by Y45, establishing a steric gate distinct from other connexins. Ca^2+^ associates with E41, stabilizing the neck via electrostatic remodeling without global conformational change. Together, these data define a dual steric-electrostatic mechanism for Cx45 regulation and provide a structural framework for isoform-specific connexin gating relevant to cardiac physiology and conduction disease.

## Introduction

Twenty-one connexin isoforms have been identified in the human genome. Together, they form a family of transmembrane proteins that assemble into gap junction channels, enabling the direct exchange of ions, metabolites, and signaling molecules between adjacent cells. Each gap junction channel is formed by the end-to-end docking of two hemichannels (connexons), each composed of six connexin subunits^1,2^. Through this intercellular communication pathway, connexins coordinate electrical and metabolic coupling across tissues, regulating synchronized contraction in the heart, neuronal activity in the brain, epithelial barrier function in the skin, and homeostasis in sensory organs such as the cochlea and retina^3–5^. Gap junction–mediated coupling is therefore essential for tissue development, physiological signaling, and maintenance of organ function. Given this central role, disruption of connexin structure or regulation is associated with a broad spectrum of human diseases. For example, mutations in GJB2 (Cx26) cause nonsyndromic hearing loss and keratitis-ichthyosis-deafness syndrome; GJA1 (Cx43) mutations are linked to oculodentodigital dysplasia; GJB1 (Cx32) mutations underlie X-linked Charcot–Marie–Tooth neuropathy; and alterations in cardiac connexins (Cx40, Cx43, Cx45) contribute to arrhythmias and conduction abnormalities^5–12^. Aberrant connexin expression or misregulation has also been implicated in cancer progression, inflammation, and impaired wound healing, underscoring the importance of understanding the structural and regulatory principles governing gap junction function^13,14^.

Connexin 45 (Cx45), encoded by the GJC1 gene, is one of the major isoforms in the cardiovascular system. It is expressed in the atrioventricular (AV) node, His bundle, and Purkinje fibers, where it contributes to the slow conduction pathways critical for normal cardiac rhythm^15–17^. Beyond the specialized conduction system, Cx45 is also expressed at lower levels in working ventricular myocytes and is upregulated in cardiomyocyte-restricted Cx43-knockout mice, where it partially compensates for the loss of Cx43 and supports residual ventricular conduction^18^. Genetic disruption of GJC1 in mice results in embryonic lethality due to cardiovascular defects, underscoring its essential role in development^19,20^. In humans, several Cx45 mutations including R75H, R184G, and M235L have been linked to cardiac conduction abnormalities, familial arrhythmias, and congenital heart disease^9,10,21–23^. Of particular interest is the R75H mutation, located near the extracellular vestibule, a region implicated in Ca^2+^-dependent gating, which is associated with atrioventricular block and arrhythmia^22–24^. Despite its physiological importance and disease relevance, the molecular basis of Cx45 channel gating and regulation has remained undefined.

Over the last decade, structural biology methods, especially cryogenic electron microscopy (cryo-EM), have revolutionized our understanding of connexin structure and function. High-resolution structures of several connexin isoforms, including Cx26, Cx46/50, Cx32, Cx36, Cx31.3, and Cx43, have revealed the overall architecture of either hemichannel or gap junction channel, as well as the arrangement of the NTD, transmembrane helices (TM1–TM4), and extracellular loops (ECL1, ECL2)^25–34^. These studies have also provided structural details of distinct functional states, from open to closed conformations, and have illuminated gating mechanisms involving primarily the NTD, bound lipids, and Ca^2+^ binding in the channel vestibules. For example, the crystal structure of Cx26 gap junction channels in the presence of Ca^2+^ revealed E42, G45 and E47 coordinating Ca^2+^ ions to generate a positive electrostatic barrier, consistent with a closed state^25^. Recent cryo-EM structures of Cx43 captured a fully closed state in which the NTDs line the pore and create seven small openings^30^, whereas Cx31.3 exhibits a semi-closed configuration with the NTD constricting the pore without closing completely at the cytoplasmic entrance^31^. However, the structural roles of the cytoplasmic intracellular loop (IL) and the flexible C-terminal domain (CTD) have remained poorly understood, limiting our understanding of how intracellular regions contribute to channel assembly and regulation.

Despite major advances in connexin structural biology, no experimental structure of Cx45 has been reported in either hemichannel (HC) or gap junction channel (GJC) form, leaving its molecular architecture and gating mechanism unresolved. It is unclear whether Cx45 employs gating features analogous to those described for other connexins, such as NTD-mediated pore constriction and extracellular Ca^2+^-dependent regulation, or whether isoform-specific structural elements fine-tune pore geometry and gating behavior within the conserved connexin framework. In particular, the gating mechanism, the structural basis of Ca^2+^ coordination, and the mechanistic implications of disease-associated mutations remain undefined.

Here, we present the high-resolution cryo-EM structures of the human Cx45 GJC in apo and Ca^2+^-bound states in addition to an E41A mutant. These structures were resolved at 2.76 Å, 2.65 Å and 3.55 Å resolution, respectively. Our study reveals that although Cx45 preserves the canonical connexin architecture, its pore constriction is defined by a ring of Y45 residues at the extracellular neck region, establishing a previously unrecognized steric gating site. The NTD, in turn, adopts a semi-closed configuration that lines the cytoplasmic vestibule without fully occluding it. In the Ca^2+^-bound state, E41 coordinates calcium ions within a conserved acidic microenvironment at the extracellular vestibule, inducing localized conformational and electrostatic remodeling without global structural rearrangements. Mutational, biochemical, and molecular dynamics analyses further support roles for Y45 in steric gating and E41 in Ca^2+^ coordination. Together, these findings provide a structural framework for understanding Cx45 gating and regulation and illustrate how conserved connexin architecture can be fine-tuned by isoform-specific pore determinants to support specialized functions.

## Results

### Cryo-EM structure of human Cx45 (GJC1) gap junction channel

The human Cx45 GJC was expressed in Expi293 cells. The purified Cx45 with a C-terminal eGFP-Strep-tag was analyzed by size exclusion chromatography, which showed two major peaks corresponding to the dodecameric GJC and the hexameric HC, respectively (**Fig. 1a, Supplementary Fig. 1**). Fractions corresponding to the GJC and HC peaks were pooled, concentrated, and used for cryo-EM grid preparation. Representative cryo-EM micrographs showed well-dispersed particles (**Fig. 1b** and **Supplementary Fig. 2a**). While particles corresponding to both GJC and HC assemblies were observed, HC particles predominantly exhibited top views, resulting in limited angular sampling that hindered high-resolution 3D reconstruction. In contrast, GJC particles assumed a broad distribution of orientations, including clear side views, enabling three-dimensional reconstruction. The 3D-classification and refinement with D6 symmetry yielded a map at 2.76 Å resolution, in which the transmembrane region was surrounded by LMNG detergent micelle density forming characteristic belt-like features (**Fig. 1c and Supplementary Fig. 2b-f**).

**Figure 1.**
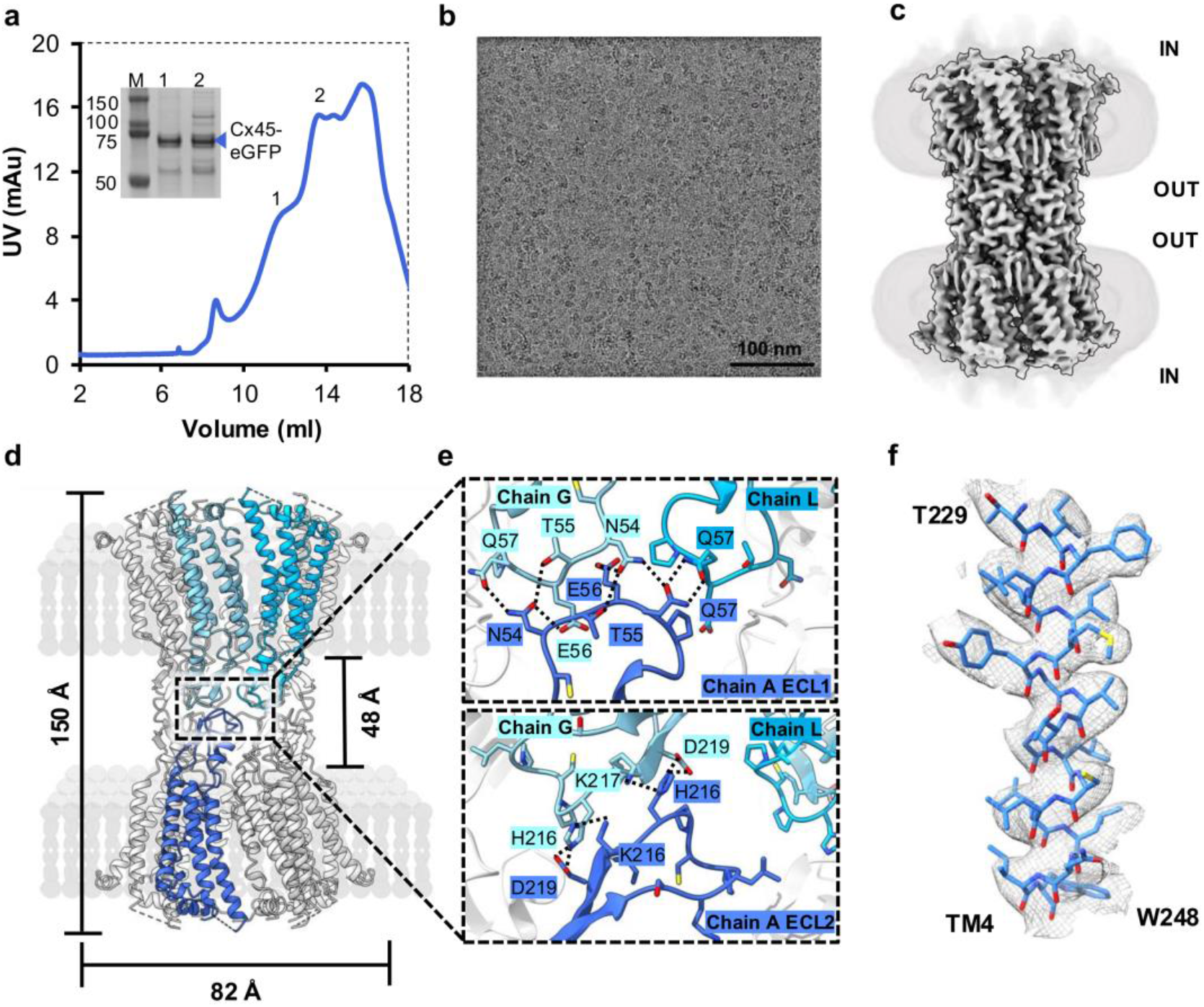
Cryo-EM structure of human Cx45 gap junction channel (GJC). **a.** Size exclusion chromatography profile of Cx45-GJC fused with eGFP. The two peaks labeled 1 and 2 correspond to GJC and HC, respectively. An SDS-PAGE analysis of the full-length Cx45-eGFP (72 kDa) protein bands and markers is shown in the inset. **b**. A representative cryo-EM micrograph of Cx45-GJC. **c**. A cryo-EM map of Cx45-GJC, with micelle belts (light gray) covering the transmembrane regions **d**. The atomic model of Cx45-GJC, fitted into the cryo-EM map, shown in a side view. Representative protomers in both hemichannels are highlighted to visualize the hemichannel docking interface. Two hemichannels (HC) dock face-to-face to form a full GJC **e**. A zoom-in view of the HC docking interface through ECL1 and 2, highlighting the hydrogen bonds and salt bridges with dotted lines. **f**. A zoom-in view showing the quality of the atomic model fit to the cryo-EM map in the TM4 region (**Supplementary Fig. 5a**).

The cryo-EM structure of human Cx45-GJC reveals a canonical dodecameric assembly composed of two hexameric HCs arranged in a head-to-head configuration across the extracellular space (**Fig. 1d**). Each protomer adopts the conserved connexin fold, comprising four transmembrane helices (TM1–TM4), two extracellular loops (ECL1 and ECL2), an intracellular loop (IL), an N-terminal domain (NTD), and a long, flexible C-terminal domain (CTD) (**Supplementary Fig. 5a-c**). The extracellular docking interface is stabilized by conserved cysteine residues: each protomer contains three cysteines in ECL1 and three in ECL2, forming three intramolecular disulfide bonds that rigidify the extracellular domains and promote precise hemichannel alignment. Docking is mediated through extensive ECL–ECL interactions between opposing hemichannels, whereby ECL1 residues from one protomer engage ECL1 residues from two protomers of the opposing hemichannel, while ECL2 forms complementary contacts with its counterpart across the interface (**Fig. 1e**). This coordinated extracellular pairing generates a continuous aqueous pore spanning the intercellular gap.

The central pore is formed by contributions from TM1 and the N-terminal helices, which line the cytoplasmic vestibule and shape the pore constriction. The NTDs adopt an orientation approximately orthogonal to the pore axis, without fully occluding the channel, consistent with a low conductance or semi-open state. The transmembrane helices pack tightly to form a membrane-spanning assembly, while extracellular domains exhibit well-resolved density, enabling an accurate model of disulfide bonds and side chains (**Fig. 1f and Supplementary Fig. 5a**). In contrast, the cytoplasmic intracellular loop (residues 104–171) and distal C-terminal domain (residues 264–396) show weak or absent density, indicating conformational flexibility. Nevertheless, our cryo-EM reconstruction reveals density corresponding to a part of the intracellular loop (IL) and proximal C-terminal domain (CTD), enabling modeling of approximately 10–20 additional CTD residues compared to previously reported connexin structures, in which the CTD is typically fully disordered. The elongated CTD approaches the IL and forms a contact near K175 (**Supplementary Fig. 5c–f**), positioning IL residues L173–M174– K175 adjacent to CTD residues R259, L262, and N263, with K175 and N263 within potential interaction distance. Although this region exhibits weaker density than the transmembrane core, the observed IL–CTD proximity suggests a conformational state in which intracellular domain coupling may contribute to channel organization within the membrane bilayer.

### Y45 defines a unique neck-gating mechanism in Cx45

Cx45 NTDs project toward the cytoplasmic entrance of the pore and adopt a conformation parallel to the membrane plane. Notably, the NTDs do not converge to occlude the channel; instead, they form a semi-closed arrangement (previously shown for Cx31.3^31^) that leaves a cytoplasmic opening of ∼18 Å in diameter. This configuration suggests that in Cx45 the NTD does not constitute the principal constriction site of the pore (**Fig. 2a**). The narrowest region of the Cx45 channel is instead located at the extracellular neck region, near the interface of TM1 and ECL1. At this position, Y45 from each of the six protomers projects toward the pore axis, forming a symmetrical ring that constricts the solvent-accessible diameter to ∼6 Å, as determined by HOLE analysis^35^. Multiple sequence alignment across human connexins reveals that Y45 is unique to Cx45; most other connexins harbor smaller residues (commonly glycine) at the equivalent position, Cx36 being a notable exception containing an aspartate (**Fig. 2b**). Comparison with previously resolved connexin structures highlights a fundamental architectural shift in the location of the gating constriction (**Fig. 2c-h**). In previously resolved connexin structures, including Cx43 GJCs in gate-covering (GCN) and pore-lining (PLN) NTD conformations, Cx26 and Cx36 GJCs in PLN conformations, and Cx31.3 and Cx32 hemichannels in semi-closed conformations, the narrowest pore region is typically defined by NTD positioning, with diameters ranging from ∼6–12 Å. In Cx45, by contrast, constriction is established at the extracellular neck by Y45, defining the dominant steric gate (**Fig. 2c**). This redistribution of structural control suggests that Cx45 employs a gating strategy distinct from other connexin isoforms. To test the structural contribution of Y45, we modeled a Y45G substitution in silico; the minimum pore diameter increased to ∼14 Å, comparable to pore dimensions observed in open or PLN states of other connexins (**Fig. 2i**). These data identify Y45 as the principal structural determinant of pore constriction in Cx45. Together, these observations support a dual-layer gating architecture in Cx45. The NTD forms a size-selective constriction at the cytoplasmic entrance, whereas Y45 defines the dominant steric gate at the extracellular neck.

**Figure 2.**
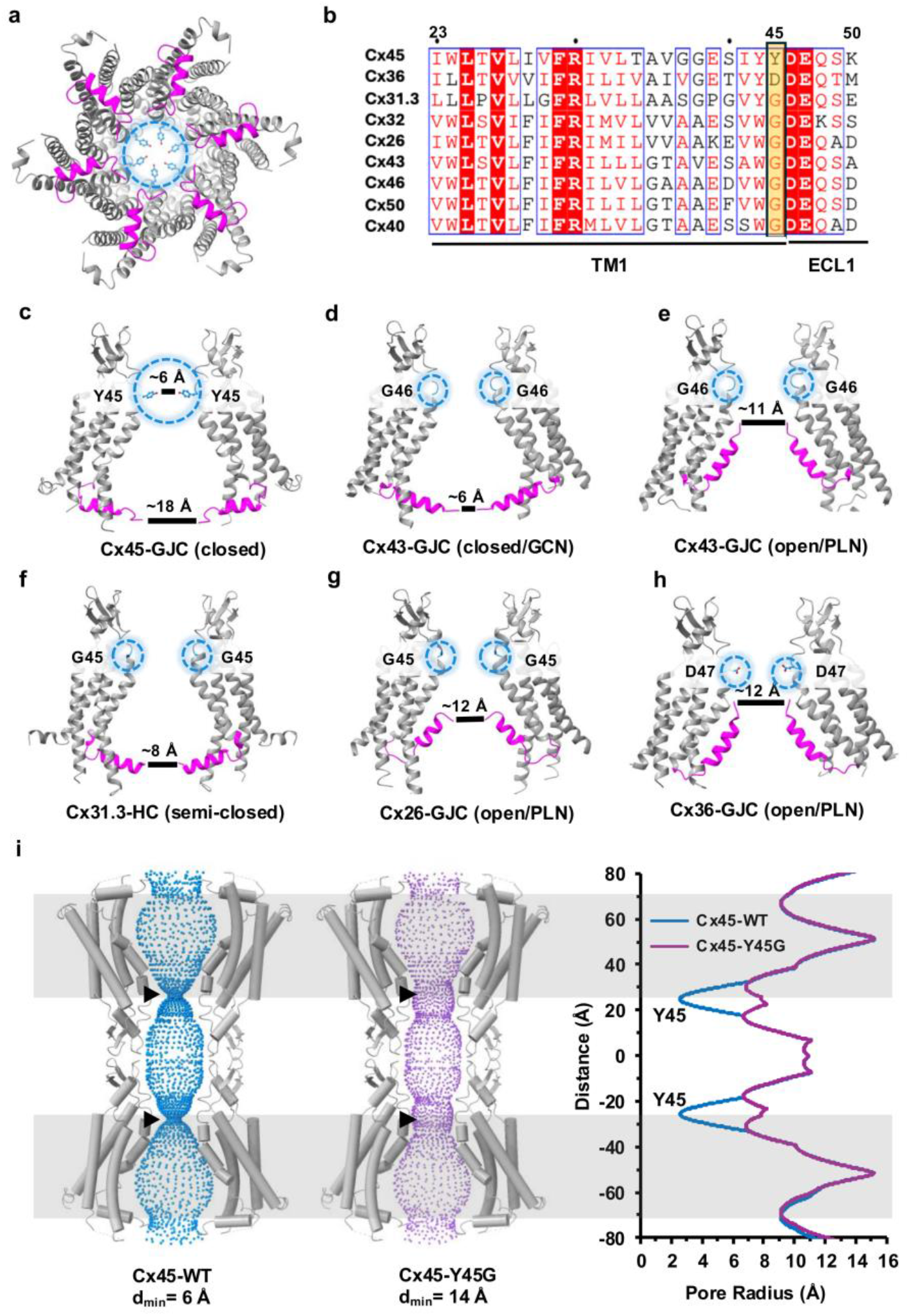
Conformation of NTD and unique pore architecture in Cx45. **a.** A cartoon representation of Cx45-GJC atomic model (top view), highlighting the conformation of NTD colored in magenta and Y45 residues (blue stick model) in the pore. **b**. A multiple protein sequence alignment of the TM1-ECL1 region in different connexins (including those with a known structure in addition to cardiac connexins), highlighting the unique Y45 residue in Cx45. **c-h**. The NTDs colored in magenta for different connexins (only two chains shown for simplifying the view). The smallest channel pore diameter is ∼ 6 Å in Cx45-GJC (**c)**, ∼6 Å in Cx43-GJC closed state (PDB: 7Z22) (**d**), ∼11 Å in Cx43-GJC open state (PDB: 8XBM) (**e)**, ∼8 Å in Cx31.3_HC semi-closed state (PDB: 6I3T) (**f)**, ∼12 Å in Cx26_GJC PLN (PDB: 2ZW3) (**g**) and ∼12 Å in Cx36_GJC PLN state (PDB: 8XGD) (**h**). The Y45 residue in Cx45 and corresponding residues (G or D) at the equivalent position in other connexins are labeled. **i**. The channel pore architecture of Cx45-GJC in WT and Y45G mutant and their representation is plotted showing the channel pore minimum diameter (d_min_) is only 6 Å (vs. 14 Å in Y45G mutant) at the Y45 site. Black arrowheads mark the Y45 constriction site in each hemichannel.

### E41 in TM1 defines a Ca^2+^-associated site at the extracellular neck

To define the structural basis of calcium regulation in Cx45, we determined the cryo-EM structure of the Cx45-GJC in the presence of Ca^2+^ (5 mM CaCl_2_) at 2.65 Å resolution (**Fig. 3a and Supplementary Fig. 3**). The Ca^2+^-bound structure closely resembles the apo state, with a Cα RMSD of 0.78 Å for the ordered transmembrane and extracellular core (TM1–TM4 and ECL1– ECL2) upon protomer superposition, indicating structural conservation of the channel scaffold with localized rearrangements near E41 and Y45. An additional density, which we attribute to Ca^2+^ ions, is observed at the channel neck region near TM1. Each Ca^2+^ ion modeled in this density is coordinated by E41 from the closest protomer together with two ordered water molecules (**Fig. 3b-c**). These six Ca^2+^ ions (per hemichannel) are positioned 3.5-4.1 Å from their closest E41 sidechain carboxylates, consistent with a loosely coordinated interaction, likely mediated by water molecules. To test whether E41 is a key coordination residue for calcium association, we generated an E41A mutant and determined its cryo-EM structure in the presence of calcium under identical conditions (**Supplementary Fig. 4)**. The E41A structure is nearly indistinguishable from the WT apo conformation and, as expected, it lacks corresponding calcium density in the neck region (**Fig. 3b**).

**Figure 3.**
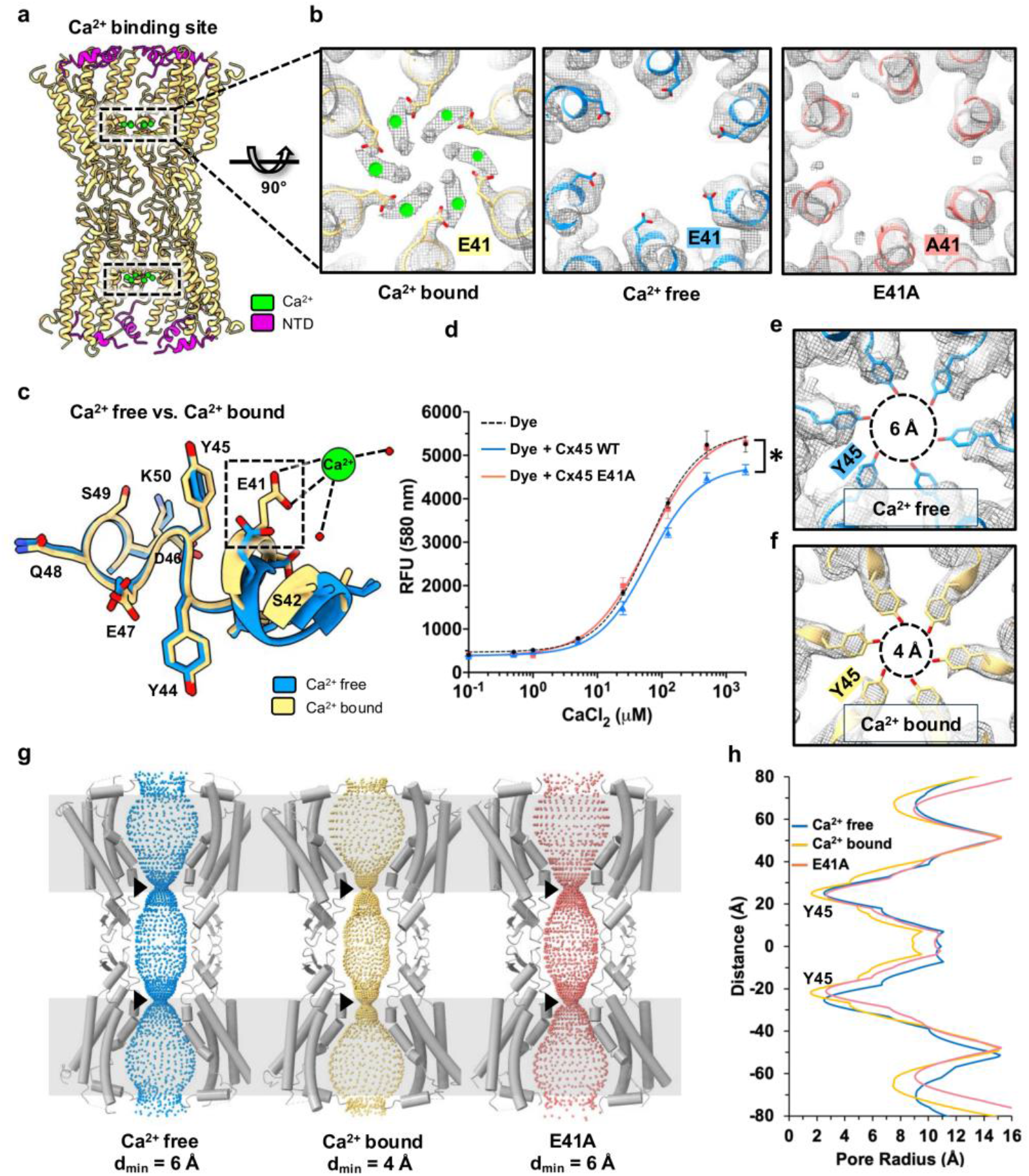
Cryo-EM structure comparison of Cx45-GJC with and without Ca^2+^. **a.** A cartoon tubular structural representation of Ca^2+^ bound Cx45-GJC, showing a Ca^2+^ binding site at the channel neck. Only four chains per hemichannel are shown for a better view of the channel pore. Ca^2+^ is shown as green spheres and NTD colored in magenta. **b**. A sliced top and zoomed in view of the cryo-EM map (in mesh) and model at the calcium binding site. The calcium bound, calcium free and E41A are shown in wheat, blue and salmon, respectively, and calcium is shown as green spheres. **c**. An overlay of TM1-ECL1 interface of Ca^2+^ bound (wheat) and Ca^2+^ free (blue) Cx45-GJC, showing the conformational rearrangement of E41 residue. Calcium is coordinated by E41 and water molecules (red spheres). **d**. A calcium competition binding assay curve of Cx45 WT and E41A mutant using RhodFF-tripotassium dye and the EC_50_ value for dye alone and with proteins are calculated from three independent protein purifications. The statistical significance marked with an asterisk represents p (WT vs. E41A) = 0.0273. **e-f**. A zoomed in view of the pore Y45 site in Ca^2+^ bound (wheat) and Ca^2+^ free (blue) Cx45-GJC. Calcium binding stabilizes the Y45 conformation and restricts the pore further. The pore d_min_ in both structures is reported. HOLE-pore analysis of three Cx45-GJC (Ca^2+^ bound (wheat) and Ca^2+^ free (blue), E41A mutant (salmon)) structures showing the pore architecture (**g**) and corresponding 1D plot (**h**) highlighting the pore constriction further in Ca^2+^ bound state, whereas E41A has a similar pore dimension as WT. The black arrowheads in panel **g** mark the Y45 constriction site in each hemichannel.

To quantify Cx45 calcium binding biochemically, we employed a RhodFF fluorescence competition assay. Dye alone yielded an EC_50_ of 57 μM (95% CI: 48–71 μM). Addition of WT Cx45 shifted the EC_50_ to 72 μM (95% CI: 64–82 μM), consistent with competition between the protein and dye for Ca^2+^. The E41A mutant, however, showed an EC_50_ of 58 μM (95% CI: 50–68 μM), significantly lower than that of WT Cx45 (extra sum-of-squares F-test, p = 0.0273) and indistinguishable from dye alone (**Fig. 3d**). These findings are consistent with loss of Ca^2+^ coordination at the E41 site, as suggested by the structural data. Structural analysis further indicates that Y45 adopts a more stabilized, inward-facing conformation in the presence of Ca^2+^, reducing the pore diameter from ∼6 Å in the apo state to ∼4 Å in the Ca^2+^-bound state (**Fig. 3e-g**). This additional constriction is absent in the E41A mutant, which structurally resembles the apo form (**Fig. 3g-h**). Together, these findings indicate that Ca^2+^ association at E41 does not drive global channel closure but instead stabilizes a locally constricted neck configuration mediated by Y45.

### Calcium binding induces an electrostatic barrier at the Cx45 channel neck

To assess how calcium binding alters the electrostatic landscape of the Cx45 pore, we performed Poisson–Boltzmann electrostatic calculations using APBS^36^ on the apo, Ca^2+^-bound, and E41A mutant structures (**Fig. 4**). In the calcium-free state, the channel neck region surrounding E41 exhibits a pronounced negative electrostatic potential, forming a deep potential well at the constriction site. Upon calcium binding, this negative potential is markedly neutralized and locally shifted toward positive values at the neck, despite the absence of large-scale conformational rearrangements in the overall structure (**Fig. 4a-b**). Cross-sectional electrostatic surface representations and one-dimensional potential profiles along the pore axis reveal a clear transition from a negative potential minimum in the apo structure (∼ −5 kT/e) to a positive electrostatic peak (∼ +15 kT/e) in the calcium-bound state at the same axial position. In contrast, the E41A mutant displays a substantially attenuated potential change, consistent with the loss of calcium coordination at this site (**Fig. 4a-c**).

**Figure 4.**
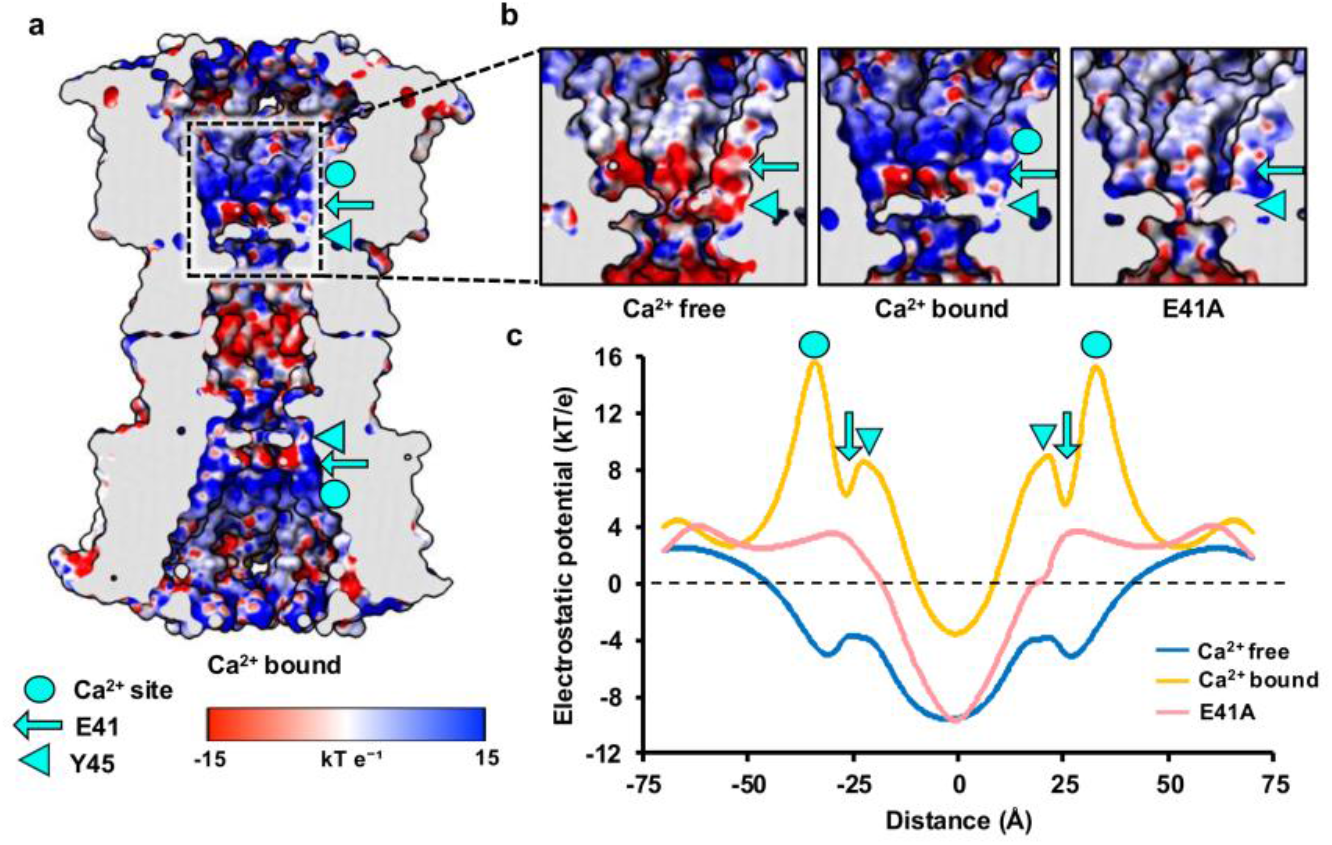
Electrostatic potential surface of Cx45-GJC. **a.** The APBS-electrostatic potential surface of Ca^2+^ bound Cx45-GJC. **b**. Zoomed in view of the Ca^2+^ binding site in Ca^2+^ free, Ca^2+^ bound and E41A structure. **c**. A 1D plot showing the APBS-electrostatic potential quantified across the channel in these Cx45-Ca^2+^ free (blue), Ca^2+^ bound (wheat) and E41A (salmon) structures. The Ca^2+^ site, E41 and Y45 are identified with symbols.

Importantly, this electrostatic remodeling occurs without major steric occlusion of the pore. Although calcium binding induces only subtle local rearrangements near E41 and a slight inward stabilization of Y45, the dominant effect is the generation of a positive electrostatic barrier at the neck. Such a charge-mediated mechanism would be expected to disfavor the permeation of positively charged ions. Together with our structural and mutational analyses, these data support a model in which calcium binding at E41 converts the narrowest part of the negatively charged vestibule into a positively biased electrostatic gate, while Y45 provides an additional steric constriction layer. This dual contribution establishes an electrostatic gating mechanism in Cx45 that operates primarily through local charge redistribution rather than global conformational change.

### Molecular dynamics simulations reveal conformational flexibility and stabilization of Y45

To investigate the structural dynamics of the extracellular pore constriction in Cx45, we performed all-atom molecular dynamics (MD) simulations of the hemichannel embedded in a lipid bilayer under both calcium-bound and apo states (see Methods). Three independent replicas were simulated for each condition, yielding a total of 3 µs of sampling per system. In these simulations, Y45 residues exhibited substantial sidechain flexibility at the pore constriction site, with the minimum distance between opposing Y45 side chains fluctuating over a broad range (∼6–10 Å) throughout the trajectories (**Fig. 5b**), reflecting continuous sampling of the pore constriction, as illustrated by representative tighter and wider conformations (**Fig. 5c**). To relate these side chain dynamics to pore architecture, we performed HOLE analysis on representative frames corresponding to tight and wide Y45 sidechain conformations. These analyses reveal pore diameters of ∼4 Å and ∼8 Å at the Y45 constriction site, respectively, demonstrating a direct correspondence between the Y45 side chain separation and the solvent-accessible pore radius (**Fig. 5d**).

**Figure 5.**
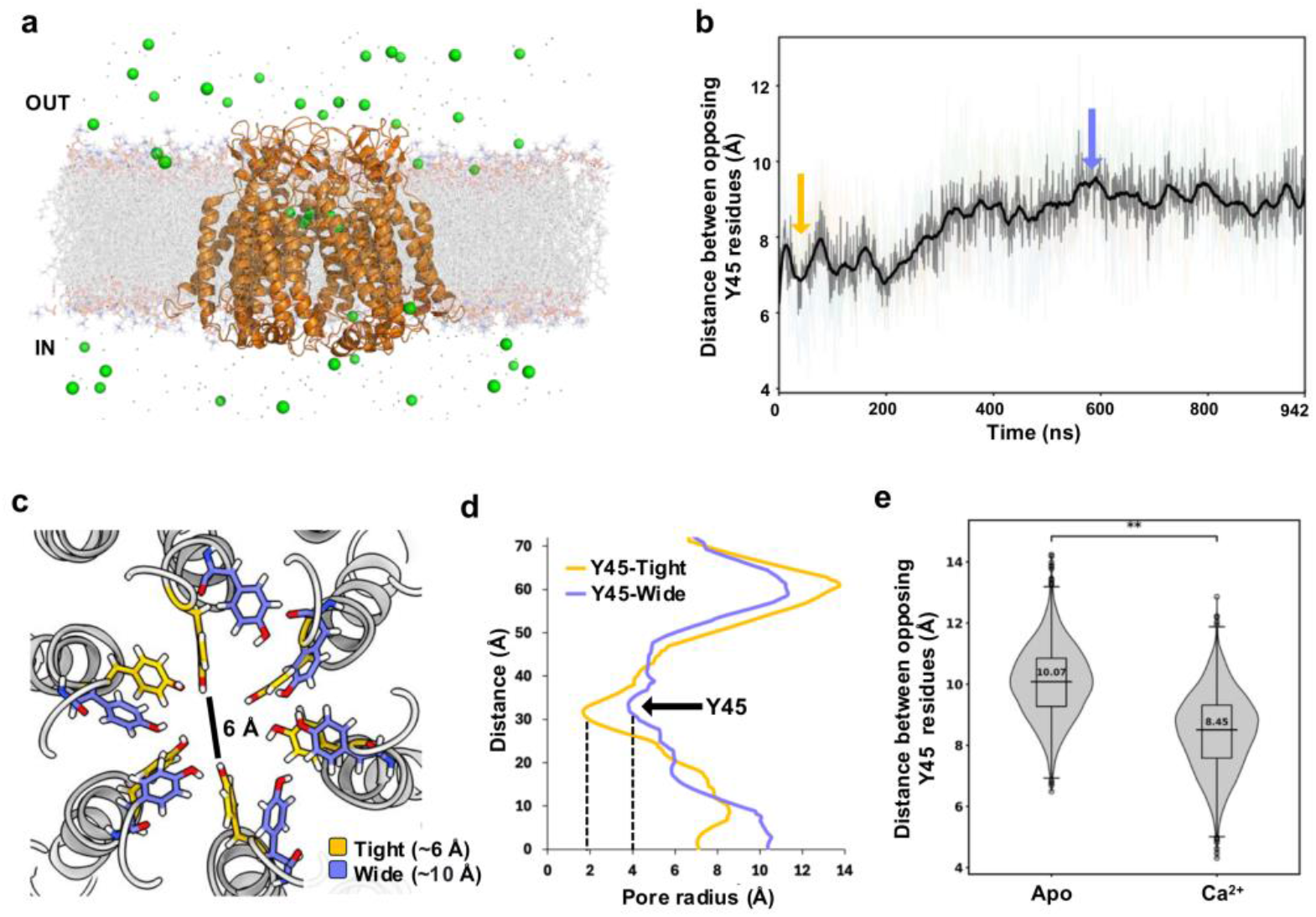
Y45 flexibility and dynamic calcium sampling at the E41 ring modulate pore constriction in Cx45. **a.** All-atom molecular dynamics simulation of the Cx45 hemichannel embedded in a lipid bilayer in the presence of Ca^2+^ ions (green spheres). The snapshot illustrates the extracellular distribution of calcium ions surrounding the channel. The sodium ions are shown as smaller spheres. **b**. Time evolution of the minimum distance between opposing Y45 residues during calcium-containing MD simulations. Individual replicate trajectories are shown as faint lines, with the mean (gray) and a smoothed mean (black) overlaid, revealing continuous fluctuations in pore constriction over a broad range (∼6–10 Å). The yellow and light blue arrows highlight the tight and wide conformation states of the Y45 sidechain during simulation. **c**. Structural representation of the Y45 constriction site illustrating the Y45 sidechain positions in tight (yellow) and wide (light blue) conformations during calcium-containing simulation. The distance between two opposing Y45 residues in tight conformation (6 Å) is marked with a black line. **d**. HOLE analysis of representative frames for the Y45 sidechain conformations showing corresponding differences in solvent-accessible pore diameter (tight = 4 Å and wide = 8 Å) at the extracellular neck. The black arrow marks the position of Y45 in both states and the dashed line defines the pore radius. **e**. Comparison of Y45 minimum distance distributions in apo (Ca^2+^-free) and calcium-containing simulations. Violin plots of Y45 minimum distance distributions indicate a clear shift toward smaller distances in the presence of calcium (mean ∼8.45 Å) relative to apo conditions (mean ∼10.07 Å; Welch’s t-test, p < 0.01, n = 3 independent trajectories per condition), demonstrating that calcium biases the pore toward more constricted conformations.

To evaluate the effect of calcium on pore architecture, we compared the distribution of opposing Y45 distances across the calcium-bound and apo conditions. Calcium shifted the system toward more constricted states, with average Y45 spacing reduced from ∼10 Å in the apo state to ∼8 Å in the calcium-bound state (**Fig. 5e**). This indicates calcium-dependent stabilization of the neck constriction. We further examined the behavior of Ca^2+^ ions near the E41 ring. In the simulations of the calcium-bound state, Ca^2+^ ions recurrently localized within coordination distance (∼3.5-4 Å) of E41 (**Supplementary Fig. 6a-b**), consistent with the binding site observed in the cryo-EM structure. These interactions were transient, suggesting that E41 serves as a flexible coordination site capable of dynamically sampling Ca^2+^ ions. In contrast, sodium ion density was elevated near E41 under apo conditions but substantially reduced in the presence of calcium (**Supplementary Fig. 6c**), indicating that Ca^2+^ may electrostatically exclude Na+ ions from the neck region. Together, these findings highlight the dynamic and flexible nature of the extracellular constriction in Cx45 and support a model in which calcium coordination at E41 stabilizes constricted Y45 conformations and contributes to electrostatic gating at the extracellular neck.

### Lipid interactions and hydrophobic pathways modulate the Cx45 pore environment

As published connexin structures display bound cholesterol and annular lipid interactions that contribute to membrane stability ^28,34^, we next examined whether similar densities are present in our Cx45 cryo-EM maps. Consistently, annular lipid or detergent densities surround the outer surface of the transmembrane helices, consistent with membrane-embedded amphipathic molecules stabilizing the channel in the bilayer environment (**Fig. 6a**). These densities are particularly prominent along the hydrophobic surfaces of TM helices in the outer leaflet, and their elongated shapes are consistent with acyl chain–like features. Within the channel interior, additional elongated densities are present in hydrophobic grooves formed between adjacent transmembrane helices (**Fig. 6b**). These densities are consistent with bound sterol molecules and were modeled as cholesterol, with two sterol molecules per protomer in the gap junction channel (**Fig. 6c–d**). The sterol rings occupy a defined hydrophobic pocket, while the hydroxyl group is oriented toward the more polar environment of the pore center, consistent with cholesterol positioning in other connexin proteins^29,30,33,34^ (**Fig. 6e**). Given that cholesteryl hemisuccinate (CHS) was present during purification, the precise sterol identity cannot be unambiguously assigned; at this resolution, only the sterol ring scaffold can be reliably modeled. Together, these sterol molecules define a continuous groove along the transmembrane region, marking a potential lipid-accessible tunnel within the channel wall that is separate from the central aqueous pore (**Fig. 6d-e**). Ordered water molecules in the extracellular vestibule contrast with the hydrophobic sterol-binding pocket, reinforcing the compartmentalization of hydrophilic and hydrophobic environments within the channel (**Fig. 6d-e**). Notably, the sterol-binding pockets are located along TM1 and TM2, near to the Y45 gating residue and the E41 Ca^2+^-coordination site. This positioning raises the possibility that bound lipids influence local helix packing and modulate the neck constriction, though whether sterol binding is required for channel stability or actively participates in gating regulation remains to be determined.

**Figure 6.**
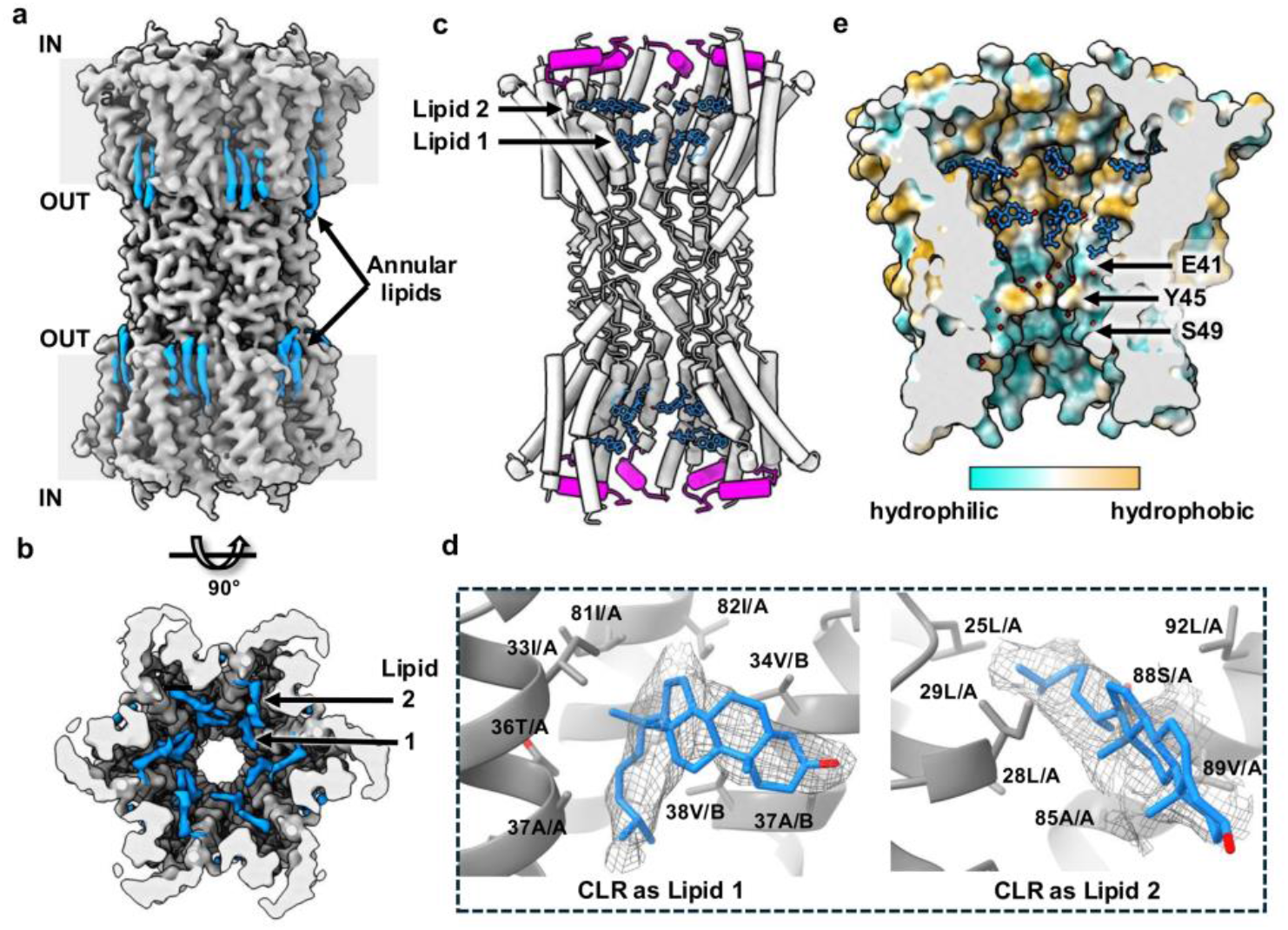
Lipid binding in Cx45-GJC. **a.** Cryo-EM map of Cx45-GJC highlighting the density of annular lipids bound on the outer surface of the transmembrane region. **b**. A sliced top view showing lipid densities inside the channel. **c**. Structural model of Cx45-GJC showing lipids 1 and 2. Cholesterol molecules are modeled in this density. Only four protomers per hemichannel are shown for clarity **d**. Density fit of cholesterol in the corresponding density, and the surrounding protein environment. **e**. Sterol molecules (cholesterol) are modeled in the hydrophobic pocket, marking a hydrophobic channel tunnel separate from the central aqueous pore. The protruding residues at the channel neck are marked and the modeled water molecules are shown as red spheres.

### Cx45 structure provides a blueprint for disease-associated mutations

Mapping the three reported Cx45 disease variants (R75H, R184G, M235L) onto the cryo-EM structure reveals two distinct structural contexts: R75 lies in the extracellular vestibule directly adjacent to the Ca^2+^-coordinating E41 and gating Y45 residues, whereas R184 and M235 reside within the transmembrane core (**Fig. 7a**). In the wild-type structure, R75 forms an electrostatic interaction network with E47 and S72 within the same protomer and with E227 from a neighboring protomer within the same hemichannel, while nearby R224 likely contributes to stabilizing the extracellular loop architecture (**Fig. 7b**). *In silico* modeling of the R75H substitution indicates disruption of this interaction network, with loss of multiple intra-protomer contacts and retention of only a single interaction between H75 and E227 (**Fig. 7c**). Given its proximity to the Ca^2+^ regulatory site, the R75H substitution would be expected to destabilize the extracellular vestibule and potentially impair both hemichannel docking and Ca^2+^-dependent gating, providing a structural rationale for the progressive conduction defects observed in patients carrying this variant^22^.

**Figure 7.**
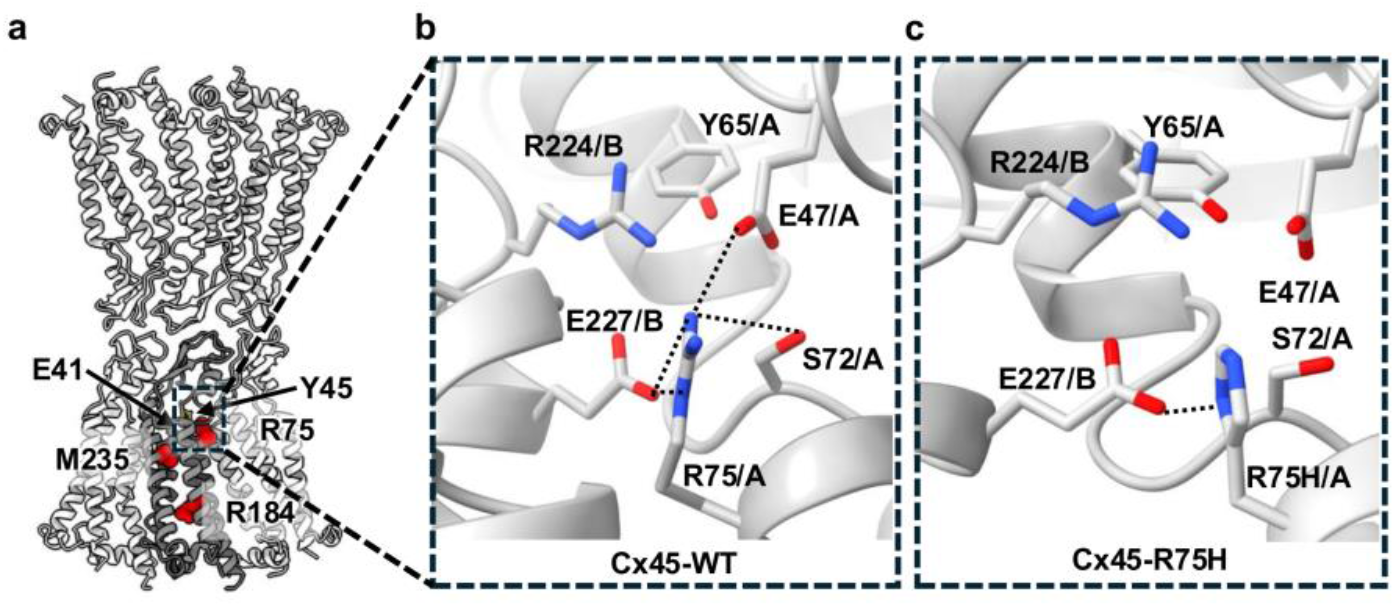
Loss of function mutation mapping in Cx45. **a.** Location of reported disease-associated mutations mapped onto the cryo-EM structure of the human Cx45 gap junction channel. Residues of R75, R184, and M235, associated with cardiac conduction defects, are shown as red spheres. Key residues involved in pore regulation and Ca^2+^ coordination (E41 and Y45) are indicated for reference. Only three protomers in each hemichannel are shown and one of the protomer is highlighted in dark gray for clarity. **b**. Structural interaction analysis of disease-associated mutation residue R75. Interactions within 3.5 Å of R75 are indicated by black dotted lines. R75 forms an interaction network involving E47 and S72 within the same protomer and E227 from the opposing protomer. R224 (chain B) is proximal to R75 (chain A) and likely contributes to the local electrostatic environment, further stabilizing the vestibule. **c**. Structural model of the R75H mutation showing disruption of the interaction network observed in the wild-type structure. In the mutant configuration, most intra-protomer interactions are lost leaving only a single contact between H75 and E227.

In contrast, R184 and M235 map to TM3 and TM4, respectively. R184 contributes a charged residue to the TM3–TM4 helix interface, and its substitution by glycine would introduce backbone flexibility at a buried position, likely destabilizing local helix packing. M235L is a more conservative substitution but may perturb a hydrophobic interhelical contact within TM4. Together, the three variants partition into two pathogenic classes — one acting on the extracellular gating machinery and the other on the integrity of the transmembrane core — illustrating how the Cx45 structure rationalizes distinct mechanisms underlying conduction disease.

## Discussion

The cryo-EM structures of human Cx45 reported here reveal a gating architecture that is fundamentally distinct from those described for other connexin isoforms. While preserving the conserved dodecameric connexin scaffold, Cx45 positions its principal permeability barrier at an extracellular neck defined by a ring of Y45 residues, rather than at the cytoplasmic NTD entrance. Ca^2+^ binds within an acidic vestibular pocket centered on E41 and remodels local electrostatics without inducing major conformational changes. Together, these observations support a dual-layer gating mechanism in which Y45-mediated steric occlusion and Ca^2+^-dependent electrostatic tuning cooperatively regulate permeation through the Cx45 channel (**Fig. 8**).

**Figure 8.**
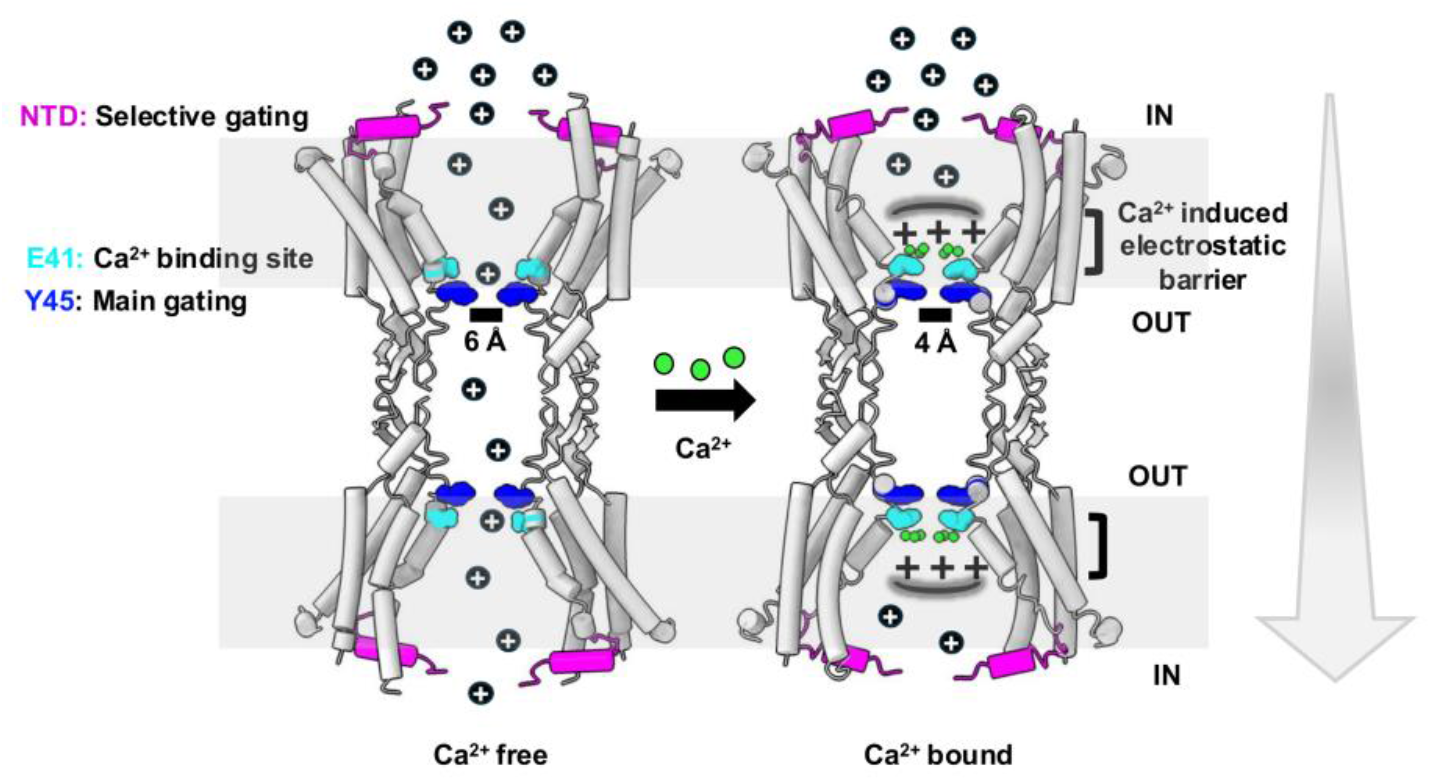
Schematic model of the proposed Cx45 gating mechanism. In the apo state, the N-terminal domains adopt a semi-closed size-selective constriction at the cytoplasmic entrance, while the narrowest region of the pore is an extracellular neck defined by Y45 (pore diameter ∼6 Å). Upon extracellular Ca^2^+ association at an acidic vestibular locus centered on E41, local electrostatics are remodeled, and the Y45 neck is stabilized in a more constricted configuration (pore diameter ∼4 Å), resulting in coupled steric and electrostatic inhibition of permeation. The NTD, E41 and Y45 are shown in magenta, cyan and blue, respectively. Bound calcium ions are shown as green spheres. Flux of cations is shown with plus signs. The direction of the gradient is illustrated with an arrow (gray).

Connexin gating is commonly mediated by NTD rearrangements at the cytoplasmic entrance, as observed in Cx26, Cx43, and related isoforms, where the NTD acts as a primary gate responsive to voltage, calcium, pH, or lipid environment^25,30,31,37^. In contrast, our structures suggest that the principal gating site in Cx45 is located at the extracellular TM1–ECL1 junction, where Y45 forms a constriction ring that narrows the pore from ∼6 Å in the apo state to ∼4 Å in the Ca^2+^-bound state. The NTD in Cx45 remains semi-closed but non-occlusive, suggesting a shift in gating control toward the extracellular domain. This neck-based gating mechanism may operate on slower timescales and respond to extracellular ionic fluctuations or hemichannel docking. The substitution of a bulky tyrosine at a position typically occupied by glycine in other isoforms exemplifies how subtle sequence variation can reprogram gating architecture within a conserved scaffold.

The ∼6-Å neck constriction observed in the apo state is just below the hydrated diameter of monovalent cations such as Na+ (∼7.2 Å) and K+ (∼6.6 Å)^38^, suggesting that such ion permeation likely involves partial dehydration and/or transient widening of the pore. Consistent with this, our MD simulations indicate that Y45 side chains exhibit conformational flexibility, allowing the pore diameter to transiently expand to ∼8 Å (**Fig. 5d**). Such dynamic fluctuations provide a plausible mechanism by which partially hydrated ions could pass through the channel while still restricting larger metabolites or second messengers. In contrast to other connexins in fully open or PLN-like states, which support the passage of molecules up to ∼1–2 kDa, Cx45 likely restricts larger solutes due to its narrower neck. This selectivity may underlie the role of Cx45 in precisely modulating ionic conductance while limiting metabolic exchange in specialized conduction tissues.

This extracellular gate is further modulated by calcium, which regulates connexins through both direct ion association and indirect interactions via calmodulin^37,39^. An E–xxx–GDE motif near the vestibule is conserved across most canonical human connexins (except Cx23), suggesting a common Ca^2+^-binding site (**Supplementary Fig. 7**). Although the central glycine is variably substituted by tyrosine in Cx45 or aspartate in Cx36, the acidic residues in the motif remain conserved. In Cx45, Ca^2+^ binds at E41, and no additional Ca^2+^ densities are observed, indicating that the neck is the primary regulatory locus. Ca^2+^ coordination generates a local electrostatic barrier that stabilizes an inward-facing Y45 configuration, and reduces the pore diameter, all without large-scale rearrangement. Such Ca^2+^-dependent electrostatic remodeling, observed previously in Cx26 and Cx31.3, appears to represent a conserved regulatory mechanism that is locally adapted through isoform-specific pore features. Beyond ionic regulation, sterol molecules bound within TM1–TM2 grooves immediately adjacent to E41 and Y45 suggest that membrane lipid composition may provide an additional modulatory layer at the same neck locus, consistent with lipid-dependent gating described recently in Cx46/50 and Cx32^33,34^.

These structural features raise important questions about how Cx45 gating influences tissue-level function, particularly in the heart. Cx45 is prominently expressed in the AV node, His bundle, and Purkinje fiber - regions responsible for conduction delay and directional signal propagation in the heart^16,17^. Although Cx45 is also expressed in working ventricular myocytes and can partially support conduction in the absence of Cx43^18^, it does not appear to be the principal mediator of ventricular impulse propagation, and its role in ventricular cell–cell coupling under normal conditions remains incompletely defined. Within the conduction system, Cx45 forms low-conductance gap junctions (∼20–40 pS) that ensure high electrical resistance and allow fine-tuned impulse timing^40,41^. Adult mice lacking Cx45 in the heart exhibit slowed AV conduction and nodal dysfunction^17^, while human Cx45 variants such as R75H cause progressive familial heart block and atrial standstill^22^. The gating architecture described here suggests an explanation for these functional roles: the extracellular neck constriction can be directly modulated by pericellular Ca^2+^ and potentially influenced by Na+ concentrations, conditions that vary during adrenergic signaling or ischemic stress^42,43^. Stabilization of the Y45 gate by Ca^2+^ coordination at E41 reduces conductance, offering a mechanism for dynamically modulating AV node impedance. Moreover, the disease-associated variant R75H is predicted to disrupt the electrostatic network surrounding the neck (**Fig. 7c**), potentially impairing channel coupling without affecting membrane localization. Two additional reported variants, R184G and M235L, map to buried positions within the transmembrane core (**Fig. 7a**) and likely act through distinct mechanisms requiring further structural and functional characterization. Together, these observations support a model in which precise structural elements in the extracellular vestibule control Cx45 gating, and small perturbations at this regulatory site can trigger pathophysiological outcomes.

Our structures provide insight into the intracellular regions of Cx45. Compared to previously reported structures of connexin isoforms, our cryo-EM reconstruction resolves an extended portion of the CTD, revealing its proximity to the IL. Specifically, the modeled residues suggest that CTD residues N263, L262, and R259 are positioned adjacent to IL residues L173-K175, forming a potential IL–CTD interface (**Supplementary Fig. 5e-f**). The CTD-IL interaction has been implicated in chemical and voltage-dependent gating in other connexins, particularly Cx43^44,45^. In addition, our recent *in situ* cryo-electron tomography (cryo-ET) study has indicated CTD-mediated lateral contacts between neighboring channels within the gap junction plaques^46^. Whether these intracellular interfaces contribute to gating regulation in Cx45 remains to be determined.

Collectively, our findings demonstrate how Cx45 achieves isoform-specific regulation by relocating the principal gating site from the cytoplasmic N-terminal entrance to an extracellular neck (**Fig. 8**). This neck-centered architecture enables graded control of permeability through combined steric (Y45) and electrostatic (Ca^2+^/E41) mechanisms. While our structures provide a high-resolution *in vitro* framework, understanding how these features operate in native tissues remains essential. Importantly, Cx45 can also assemble into heteromeric and heterotypic gap junction channels with other cardiac connexins such as Cx43 and Cx40^16,47^. How the extracellular neck-centered gating mechanism identified here is incorporated or reshaped within such heteromeric assemblies remains unknown and represents a key unresolved question in cardiac gap junction biology. *In situ* cryo-ET of Cx45-containing gap junctions, as well as defined heteromeric channel systems, will be critical to define how these structural features operate in the native cellular context. Complementary functional and electrophysiological assays of targeted mutants (e.g., Y45G, E41A, R75H, or IL–CTD interface disruptors) will further test their contribution to cardiac conduction and disease.

## Methods

### Expression and purification of human Cx45 channel

Human full-length Cx45 fused to a C-terminal eGFP and twin-Strep tag was cloned into the pHL-sec vector^48^ and transiently transfected into Expi293 cells, which were cultured at 37 °C with 5% CO_2_ for 72 h. Cells were collected, washed with 1× PBS, and resuspended in low-salt buffer (20 mM Tris pH 8.0, 150 mM NaCl, 1 mM EDTA, 1 mM EGTA, 2 mM β-mercaptoethanol, 10% glycerol) supplemented with 50 μg/mL nuclease (Thermo Fisher Scientific) and protease inhibitor cocktail (Thermo Fisher Scientific), followed by Dounce homogenization. Lysates were subjected to ultracentrifugation at 180000 g for 1 hour at 4 °C. Pellets were resuspended in high-salt buffer (20 mM Tris pH 8.0, 500 mM NaCl, 1 mM EDTA, 1 mM EGTA, 2 mM β-mercaptoethanol, 10% glycerol) and centrifuged again under the same conditions. The resulting membrane pellets were flash-frozen and stored at −80 °C. For purification, membranes were resuspended in solubilization buffer containing 20 mM Tris pH 8.0, 150 mM NaCl, 1 mM EDTA, 1 mM EGTA, 2 mM β-mercaptoethanol, 1% (w/v) LMNG (Anatrace), 0.1% (w/v) CHS (Anatrace) and solubilized for 2 hour at 4 °C, followed by ultracentrifugation to remove insoluble material. The supernatant was applied to Strep-Tactin Sepharose resin (IBA Lifesciences), washed with purification buffer (20 mM Tris pH 8.0, 150 mM NaCl, 1 mM EDTA, 1 mM EGTA, 2 mM β-mercaptoethanol, 0.005% LMNG and 0.0005% CHS) (10 column volumes), and eluted with the same buffer supplemented with 10 mM desthiobiotin (IBA Lifesciences). Eluted protein was immediately concentrated using a 100 kDa MWCO centrifugal concentrator (Merck Milipore Ltd.) and subjected to size-exclusion chromatography on a Superose 6 Increase 10/300 GL column (Cytiva) connected to an ÄKTA Go system in the same purification buffer. Fractions corresponding to Cx45 were identified by SDS– PAGE, pooled, and concentrated to ∼1 mg/mL. For Ca^2+^-bound structural analysis of Cx45 WT as well as E41A mutant, EDTA and EGTA were omitted after the solubilization step and 5 mM CaCl_2_ was included in the purification buffer. Similarly, the Cx45 WT and E41A proteins were purified in buffers lacking EDTA and EGTA during the Strep-Tactin affinity chromatography and SEC step for the calcium-binding assay.

### Cryo-EM specimen preparation and data collection

Freshly purified Cx45 protein was applied to glow-discharged UltraAuFoil R1.2/1.3 300-mesh gold grids (Quantifoil) and vitrified in liquid ethane using a Vitrobot Mark IV (Thermo Fisher Scientific) at 4 °C and 95% humidity. Grids were initially screened on a Talos Arctica transmission electron microscope operated at 200 kV. The apo Cx45 dataset was collected on a Titan Krios transmission electron microscope (Thermo Fisher Scientific) operated at 300 kV and equipped with a Gatan K3 BioQuantum direct electron detector and a 20-eV energy filter slit. Movies were recorded at a nominal magnification of 105,000×, corresponding to a calibrated pixel size of 0.83 Å, over a defocus range of 0.6 to 2.0 µm underfocus. Data were acquired with an exposure time of 1.8 s, fractionated into 45 frames, resulting in a total accumulated dose of ∼48 e−/Å^2^. For the Ca^2+^-bound and E41A datasets, data were collected on a Titan Krios (300 kV) equipped with a Falcon 4i detector and a Selectris energy filter (Thermo Fisher Scientific). Images were recorded at a nominal magnification of 165,000×, corresponding to a calibrated pixel size of 0.70 Å, with a 10 eV energy filter slit and a defocus range of 0.6 to 2.0 µm underfocus. The Ca^2+^-bound dataset was recorded with an exposure time of 3.0 s and fractionated into 51 frames, yielding a total electron exposure of 47 e−/Å^2^. The E41A dataset was collected under identical optical conditions with an exposure time of 3.1 s, resulting in a total electron exposure of 50 e−/Å^2^.

### Image processing and reconstruction

All cryo-EM datasets were processed using cryoSPARC v4.7.0^49^. The image processing and reconstruction workflows are shown in **Supplementary Fig. 2-4**. Dose-fractionated movies were motion-corrected and subjected to patch-based CTF estimation. Initial particle picking was performed using a combination of blob and manual picking to generate templates for template-based picking. Following preliminary 2D classification and curation, high-quality particles in a subset of 100 micrographs were used to train a model, which was subsequently applied for automated particle picking in Topaz^50^. Extracted particles were subjected to multiple rounds of 2D classification to remove contaminants and poorly aligned particles. Remaining particles were further cleaned using ab initio reconstruction followed by iterative heterogeneous refinement without applied symmetry to eliminate low quality particles prior to high-resolution refinement.

For the Cx45 apo dataset, the curated particle set (24,240 particles) was subjected to 3D classification without alignment to assess conformational heterogeneity. Two classes displayed poorly resolved N-terminal domain (NTD) density, whereas one class (8,690 particles) exhibited well-defined NTD features and was selected for further refinement. This homogeneous subset was refined using non-uniform refinement with D6 symmetry imposed, yielding a final reconstruction at 2.76 Å resolution based on the Fourier shell correlation (FSC) 0.143 criterion.

For the Cx45 Ca^2+^-bound dataset, particle processing was processed as described above unless stated otherwise. Following curation, non-uniform refinement with D6 symmetry initially yielded a reconstruction at 2.89 Å resolution from 15,173 particles. Enabling CTF refinement during subsequent refinement improved the final resolution to 2.65 Å. Similarly, for the Cx45 E41A dataset, the curated particle set (11,966 particles) was refined using non-uniform refinement with D6 symmetry imposed, yielding a final reconstruction at 3.55 Å resolution.

For all datasets, map sharpening was performed using automated B-factor estimation in cryoSPARC. Detergent micelle density surrounding the transmembrane region was reduced for visualization using the LocScale2 plugin in UCSF ChimeraX^51,52^.

### Model building and refinement

Model building was performed using the cryo-EM density maps for each dataset. For the Cx45 apo structure, an AlphaFold2 model corresponding to the ordered transmembrane and extracellular regions of human Cx45 was used as an initial template^53^. The model was rigid-body fitted into the cryo-EM map and further adjusted using ISOLDE in UCSF ChimeraX to optimize local geometry and improve agreement with the density^54^. The ISOLDE-refined model was subjected to iterative real-space refinement in Phenix (v1.21.2-5419-000) against the cryo-EM map^55^. Refinement included secondary structure restraints and non-crystallographic symmetry (NCS) constraints. Individual atomic displacement parameters (ADPs) were refined isotropically. Iterative cycles of manual model correction and geometry inspection were performed in Coot^56^ followed by a final round of refinement in Phenix.

In the apo structure, well-defined non-protein densities within hydrophobic grooves of the transmembrane region were interpreted as cholesterol-like molecules and thus modeled as cholesterol (CLR). A total of 24 CLR molecules were modeled. Ordered spherical densities within the extracellular vestibule and neck region, consistent with hydrogen-bonding geometry, were modeled as waters. For the Ca^2+^-bound structure, the refined apo model was used as the starting model for real-space refinement in Phenix^55^. Additional densities observed in the extracellular neck region were modeled as Ca^2+^ ions, with one ion per protomer (six per HC, twelve per GJC), based on coordination geometry and consistent density features. Calcium ions were refined with appropriate geometry restraints. Similarly, the E41A mutant structure was refined starting from the apo model, and an alanine residue was built at position 41, consistent with the density. The first N-terminal methionine residue was not modeled in any structure due to the absence of corresponding density. The N-terminal domain exhibited lower local resolution in all datasets, consistent with conformational flexibility observed during 3D classification and local resolution analysis. Model validation was performed using MolProbity within Phenix^57^. Refinement statistics are summarized in **Table 1**, and the model–map FSC curves for all three structures are shown in **Supplementary Fig. 5b**. Structural analysis and visualization were performed using UCSF ChimeraX unless otherwise stated^51^.

**Table 1.** Cryo-EM data collection, refinement and validation statistics.

|  | Cx45-GJC apo | Cx45-GJC Ca<sup>2+</sup><br>bound | Cx45-GJC<br>E41A |
| --- | --- | --- | --- |
| **Data collection and processing** |  |  |  |
| Instrument | FEI Titan Krios<br>(Gatan K3<br>BioQuantum) | FEI Titan Krios<br>(Falcon 4i) | FEI Titan Krios<br>(Falcon 4i) |
| Magnification | 105000 | 165000 | 165000 |
| Voltage (kV) | 300 | 300 | 300 |
| Electron exposure (e-/Å<sup>2</sup>) | 48 | 47.4 | 50 |
| Defocus range (μm) | 0.6 – 2.0 | 0.6 -2.0 | 0.6 – 2.0 |
| Pixel size (Å) | 0.828 | 0.70 | 0.70 |
| Symmetry imposed | D6 | D6 | D6 |
| Number of particles | 8690 | 15173 | 11966 |
| Map resolution (Å) | 2.76 | 2.65 | 3.55 |
| FSC 0.143 |  |  |  |
| Map local resolution range (Å) | 2.51 – 6.20 | 2.50 – 6.15 | 3.10 – 7.25 |
| **Refinement** |  |  |  |
| Map sharpening *B* factor (Å<sup>2</sup>) | -55 | -27.8 | -62.5 |
| Initial model used | AlphaFold model | Cx45-GJC apo | Cx45-GJC apo |
| Model resolution (Å) |  |  |  |
| FSC 0.143 | 2.7 | 2.6 | 3.5 |
| FSC 0.5 | 3.1 | 3.5 | 3.9 |
| Model composition |  |  |  |
| Non-hydrogen atoms | 19308 | 17256 | 17280 |
| Protein residues | 2328 | 2244 | 2208 |
| Ligands | 24 (CLR) | 12 (Ca) | - |
| *B* factors (Å<sup>2</sup>) |  |  |  |
| Protein | 95 | 82 | 53 |
| Ligand | 123 | 93 | - |
| R.m.s. deviations |  |  |  |
| Bond lengths (Å) | 0.002 | 0.002 | 0.002 |
| Bond angles (°) | 0.588 | 0.477 | 0.475 |
| Map CC (mask) | 0.84 | 0.76 | 0.72 |
| Validation |  |  |  |
| MolProbity score | 1.7 | 2.10 | 2.23 |
| Clashscore | 6.99 | 7.76 | 7.81 |
| Poor rotamers (%) | 0.6 | 1.4 | 2.60 |
| Ramachandran plot |  |  |  |
| Favored (%) | 95.8 | 96.2 | 93.9 |
| Allowed (%) | 4.2 | 3.8 | 6.1 |
| Disallowed (%) | 0 | 0 | 0 |

### Pore radius analysis

Pore dimensions were calculated using the program HOLE (v2.0)^35^. The refined atomic models were used as inputs, and the pore axis was defined along the six-fold symmetry axis passing through the channel center. The starting point and vector defining the pore axis were determined based on the central cavity coordinates. A sampling interval of 0.25 Å was used with a maximum radius cutoff of 20 Å and Bondi atomic radii. Output radius profiles were extracted from the HOLE log files and plotted as a function of distance along the pore axis. Surface representations of the pore were visualized in ChimeraX. For the Y45G mutant, tyrosine was substituted with glycine in ChimeraX, and the local environment surrounding position 45 was refined using ISOLDE to optimize the geometry while maintaining overall structural restraints. The refined model was subsequently subjected to identical HOLE parameters for comparative pore profiling.

### Electrostatic potential calculations

Electrostatic potentials were calculated by solving the linearized Poisson–Boltzmann equation using APBS^36^. Water molecules and non-relevant ligands were removed from all structures; Ca^2+^ ions were retained in the Ca^2+^-bound model. PQR files and APBS input parameters were generated using the APBS web server^36^. Calculations were performed using a protein dielectric constant of 2.0 and a solvent dielectric constant of 78.54 at 298.15 K, with an ionic strength of 150 mM monovalent salt. Electrostatic potential maps were generated in OpenDX (.dx) format. Electrostatic potentials were visualized in UCSF ChimeraX by mapping the potential onto the solvent-accessible surface of the protein and colored using a red–blue palette (−15 to +15 kT/e). Identical visualization parameters were applied to the apo, Ca^2+^-bound, and E41A models to enable direct comparison. To quantify electrostatic changes along the pore axis, potentials were sampled from the APBS maps along a line parallel to the pore axis at a fixed coordinate corresponding to the channel center. Electrostatic potential values were plotted as a function of distance along the pore axis (kT/e) using a custom python script (plot_apbs_1d.py), which performs trilinear interpolation of the APBS grids, and identical sampling coordinates were used for all datasets.

### Calcium binding assay

Calcium-binding assays were performed using the fluorescent low-affinity calcium indicator RhodFF tripotassium salt (AAT-Bioquest)^58,59^. Cx45 protein (WT or E41A) was purified in buffer containing 20 mM Tris (pH 8.0), 150 mM NaCl, 0.005% LMNG, and 0.0005% CHS. EDTA and EGTA were included during membrane preparation and solubilization to remove residual metal ions but were omitted during final purification and fluorescence measurements to prevent Ca^2+^ chelation and ensure accurate assessment of calcium binding. Assays were conducted in 96-well plates using an EnSpire multimode plate reader (PerkinElmer). RhodFF was used at a final concentration of 100 nM, and Cx45 was added at 1 µM where indicated. Calcium chloride was titrated over a range of 0–2000 µM (0, 0.1, 0.5, 1, 5, 25, 125, 500, and 2000 µM). Fluorescence emission was recorded at 580 nm following excitation at 552 nm using top-read mode. Fluorescence values were analyzed in GraphPad Prism (V10.6.1) using nonlinear regression (log [Ca^2+^] vs response, four-parameter logistic model with variable slope)^60^. Apparent half maximal effective concentration (EC_50_) values are reported with 95% confidence intervals. Statistical comparison between WT and E41A titration curves was performed using an extra sum-of-squares F test^61^. Experiments were performed using three independent protein purifications.

### Multiple sequence alignment

Full-length human connexin protein sequences were retrieved from UniProt and aligned using ClustalW^62^ with default parameters. The TM1–ECL1 region was analyzed to assess conservation of residues. Alignment, visualization, and annotation were generated using the ESPript 3.0 web server^63^.

### Atomistic molecular dynamics simulations

A connexin hemichannel from the cryo-EM structure of the Cx45 gap junction channel was embedded in a mixed lipid bilayer composed of POPC and cholesterol at a 70:30 molar ratio using the CHARMM-GUI membrane builder workflow^64^. The system was parameterized using the CHARMM36m force field for proteins and lipids, with TIP3P water and compatible ion parameters, which together provide improved sampling of backbone and sidechain conformations in membrane proteins. Two simulation conditions were prepared: (i) a calcium-containing system with 5 mM CaCl_2_, where calcium ions were initially placed at experimentally observed binding sites and allowed to diffuse freely during the simulations without positional restraints, and (ii) a calcium-free (apo) system without added Ca^2+^. In both setups, the ionic strength was adjusted to approximately 150 mM (NaCl) to approximate physiological conditions. All simulations were performed using GROMACS 2022^65^. Each system was first energy minimized to remove steric clashes and then equilibrated in a multi-step protocol using the NVT ensemble. Early equilibration steps employed a 1 fs timestep with positional restraints on the protein backbone and lipid heavy atoms, which were gradually released. Production simulations were performed in the NPT ensemble using a 2 fs timestep, with three independent 1000-ns replicas conducted per condition, yielding a total of approximately 3 μs of sampling for each system. All bonds involving hydrogen atoms were constrained using the LINCS algorithm^66^, enabling the use of a larger timestep. Long-range electrostatic interactions were treated using the particle mesh Ewald (PME) method with a 1.1 nm real-space cutoff^67^, and van der Waals interactions were smoothly switched off between 1.0 and 1.2 nm using a force-based switching function. Temperature and pressure were maintained at 300 K and 1 bar using the velocity-rescaling thermostat^68^ and Parrinello–Rahman barostat^69^, respectively. Trajectory analyses were performed using MDAnalysis^70^. Distances between opposing Y45 residues were calculated as the minimum distance between sidechain heavy atoms across the pore. Sodium ion density profiles along the membrane normal (Z-axis) were computed using the GROMACS density tool by binning ion positions along the Z-axis, averaging over the trajectory, and normalizing by the bin volume. Profiles were averaged across independent replicas to assess ion distribution near the pore neck.

### AI-assisted technologies

During the preparation of this manuscript, ChatGPT (OpenAI) was used to assist with code refinement. Both ChatGPT and Claude.ai (Anthropic) were used for editing the text for clarity. Authors take full responsibility of the contents.

## Code availability

The custom python script used to extract electrostatic potential values along the pore axis from APBS-generated maps and generate one-dimensional electrostatic potential profiles is available at Laboratory of Structural Biology GitHub (https://github.com/LSB-Helsinki/scripts/). All other analyses were performed using standard tools as described in the Methods.

## Acknowledgements

We thank Xun Lu, Maryna Green, Tuomas Niemi-Aro and Behnam Lak for technical assistance. This study was funded by grants from the Sigrid Jusélius Foundation (to J.T.H and to V.S.). We acknowledge CSC–IT Center for Science, Finland, for computational resources. We acknowledge the facilities and expertise of the HiLIFE cryo-EM unit at the University of Helsinki (a member of Instruct-ERIC Centre Finland, FINStruct, and Biocenter Finland) and SciLifeLab cryo-EM facility (Stockholm and Umeå, Sweden).

## Author Contributions

Investigation: S.K.S.T. and E.E. Data Curation: S.K.S.T., E.E., E.-P.K. and J.T.H. Methodology: S.K.S.T., E.E. and A.A. Formal Analysis: S.K.S.T., E.E., E.-P.K., V.S. and J.T.H. Visualization: S.K.S.T., A.K. and A.A. Conceptualization: S.K.S.T. and J.T.H. Supervision: V.S. and J.T.H. Project Administration: J.T.H. Writing – Original Draft: S.K.S.T. and J.T.H. Writing – Review & Editing: all authors. Funding Acquisition: V.S. and J.T.H.

## Supplementary Information

A supplementary information file is available for this manuscript.

