## supplementary file for "Gating Mechanism of the Human Connexin 45 Gap Junction Channel"

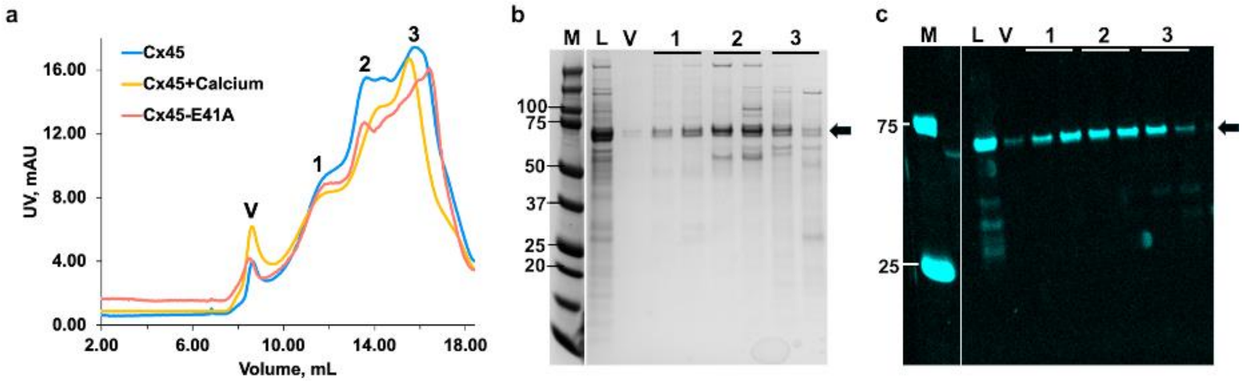


**Supplementary figure 1. Purification of Cx45. a.** An overlayed SEC purification profile of Cx45 fused with eGFP at C-terminus samples, purified in LMNG. The peak V, 1, 2, 3 corresponds to peak for void, gap junction channel, hemi channel and Cx45-impurities/truncation mixed respectively. A SDS-PAGE with Coomassie staining (**b**) and SDS-In-gel fluorescence (**c**) analysis of protein samples corresponding to each peak. The standard protein marker (M) bands, sample loaded to SEC column (L), other lanes corresponding to peaks in SEC, and protein bands corresponding to Cx45 fused with eGFP (72 kDa) are marked.

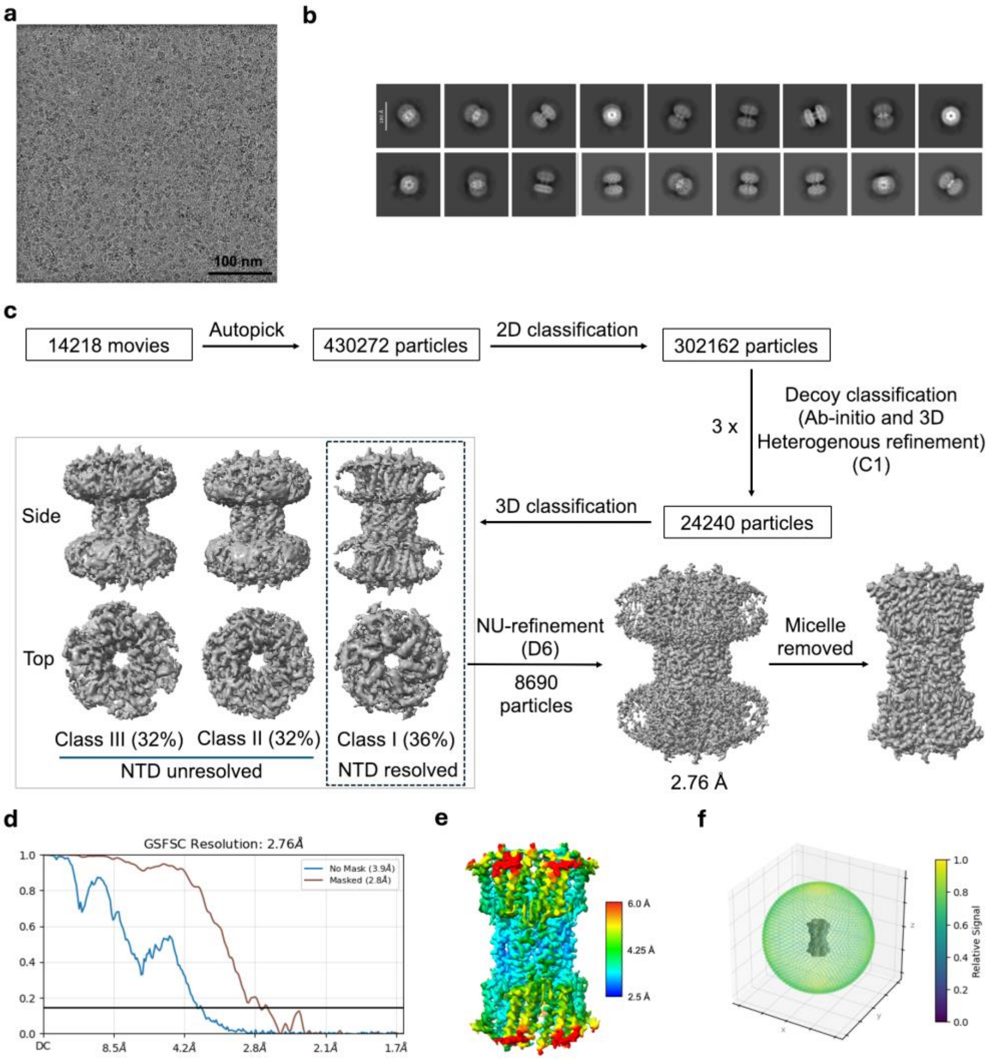


**Supplementary figure 2. Cryo-EM image processing workflow for the Cx45-apo dataset. a.** Representative cryo-EM micrograph of Cx45 gap junction channels recorded at a calibrated pixel size of 0.828 Å and a total accumulated dose of 48 $e^{-}/Å^{2}$. Scale bar, 100 nm. **b.** Representative reference-free 2D class averages following particle curation. Scale bar, 150 Å. **c.** Image processing workflow done in cryoSPARC v4.7.0. The pipeline includes motion correction, patch CTF estimation, blob/manual picking for template generation, Topaz-based particle picking, iterative 2D classification, decoy removal using *ab initio* reconstruction and heterogeneous refinement (C1 symmetry), and final non-uniform refinement with D6 symmetry imposed. Representative 3D maps at key stages of the workflow are shown. **d.** Gold-standard Fourier shell correlation (FSC) curve for the final reconstruction. The global resolution was determined to be 2.76 Å using the FSC 0.143 criterion. **e.** Local resolution estimation of the final cryo-EM reconstruction calculated in cryoSPARC and displayed on the map surface. **f.** Angular distribution of particle orientations contributing to the final reconstruction.

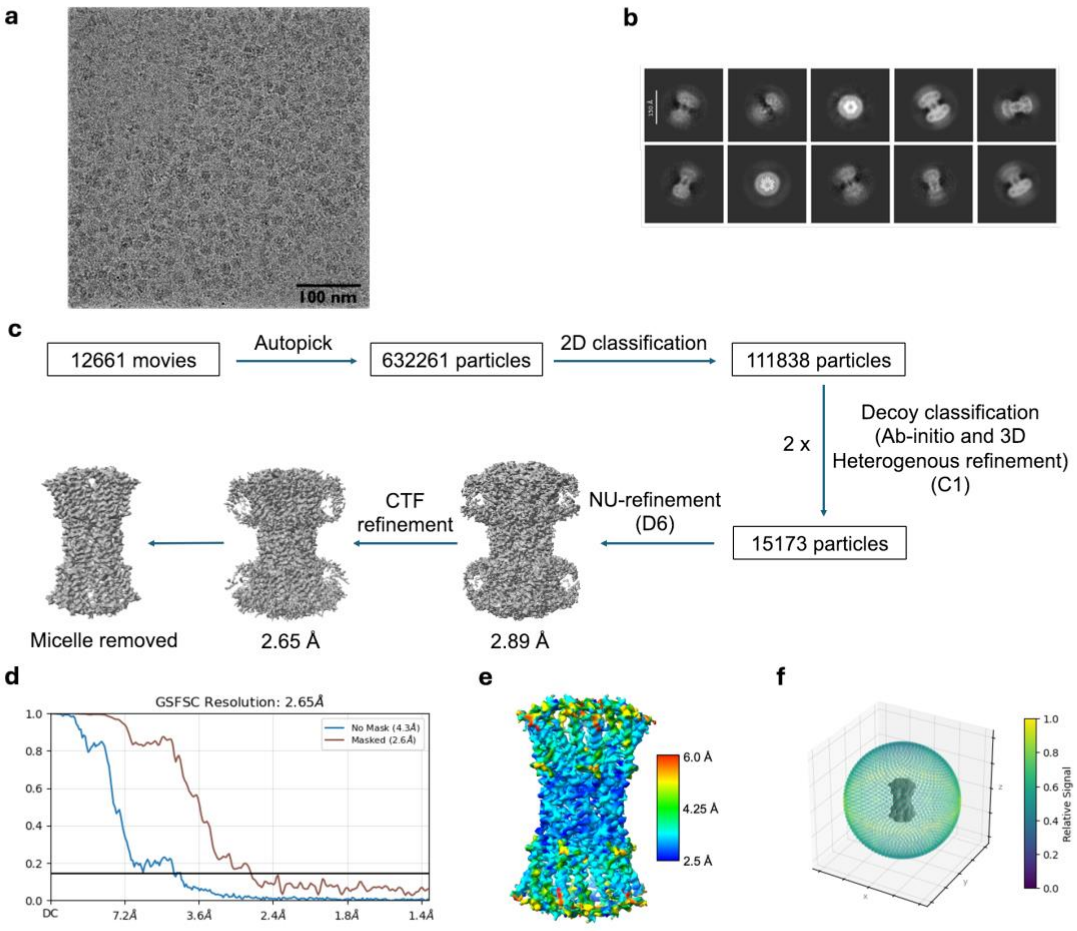


**Supplementary figure 3. Cryo-EM image processing workflow for the Cx45 $Ca^{2+}$-bound dataset. a.** Representative cryo-EM micrograph of the Cx45 $Ca^{2+}$-bound sample recorded at a calibrated pixel size of 0.70 Å with a total accumulated dose of 47.4 $e^{-}/Å^{2}$. Scale bar, 100 nm. **b.** Representative reference-free 2D class averages after particle curation, displaying well-defined views of the gap junction channel. Scale bar, 150 Å. **c.** Image processing workflow implemented in cryoSPARCv4.7.0. The pipeline included motion correction, patch CTF estimation, blob/manual picking for template generation, Topaz-based particle picking, iterative 2D classification, decoy removal using *ab initio* reconstruction and heterogeneous refinement (C1 symmetry), followed by non-uniform refinement with D6 symmetry imposed. Representative 3D reconstructions at key stages are shown. **d.** Gold-standard Fourier shell correlation (FSC) curve of the final reconstruction. The global resolution was determined to be 2.65 Å using the FSC 0.143 criterion. **e.** Local resolution estimation calculated in cryoSPARC and mapped onto the density surface of the final reconstruction. **f.** Angular distribution of particle orientations contributing to the final 3D reconstruction.

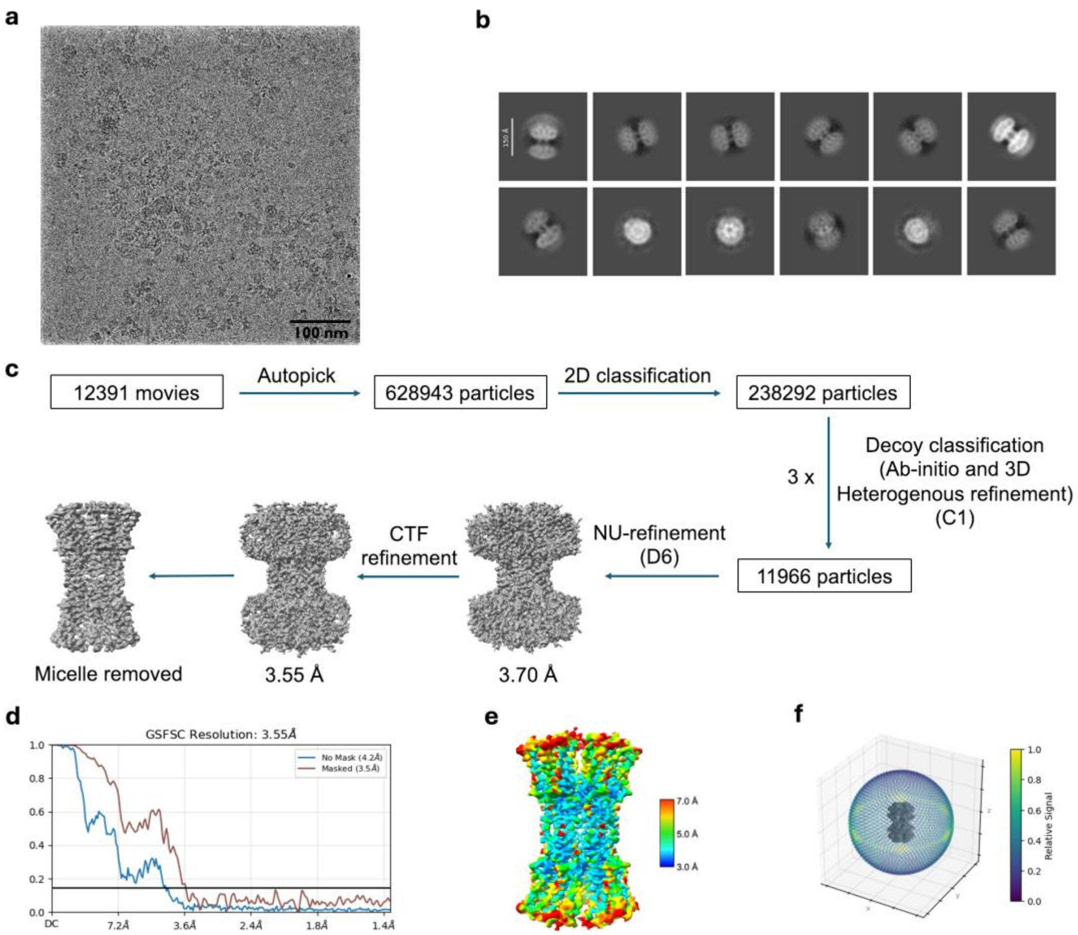


**Supplementary figure 4. Cryo-EM image processing workflow for the Cx45 E41A dataset. a.** Representative cryo-EM micrograph of the Cx45 E41A sample recorded at a calibrated pixel size of 0.70 Å and a total accumulated dose of 50 $e^-/Å^2$. Scale bar, 100 nm. **b.** Representative reference-free 2D class averages following particle curation, displaying characteristic gap junction channel views. Scale bar, 100 Å. **c.** Image processing workflow performed in cryoSPARC v4.7.0. The pipeline included motion correction, patch CTF estimation, blob/manual picking for template generation, Topaz-based particle picking, iterative 2D classification, and decoy removal using *ab initio* reconstruction and heterogeneous refinement in C1 symmetry, followed by non-uniform refinement with D6 symmetry imposed. Representative 3D reconstructions at key stages are shown. **d.** Gold-standard Fourier shell correlation (FSC) curve of the final reconstruction. The global resolution was determined to be 3.55 Å using the FSC 0.143 criterion. **e.** Local resolution estimation calculated in cryoSPARC and mapped onto the density surface of the final reconstruction. **f.** Angular distribution of particle orientations contributing to the final 3D reconstruction.

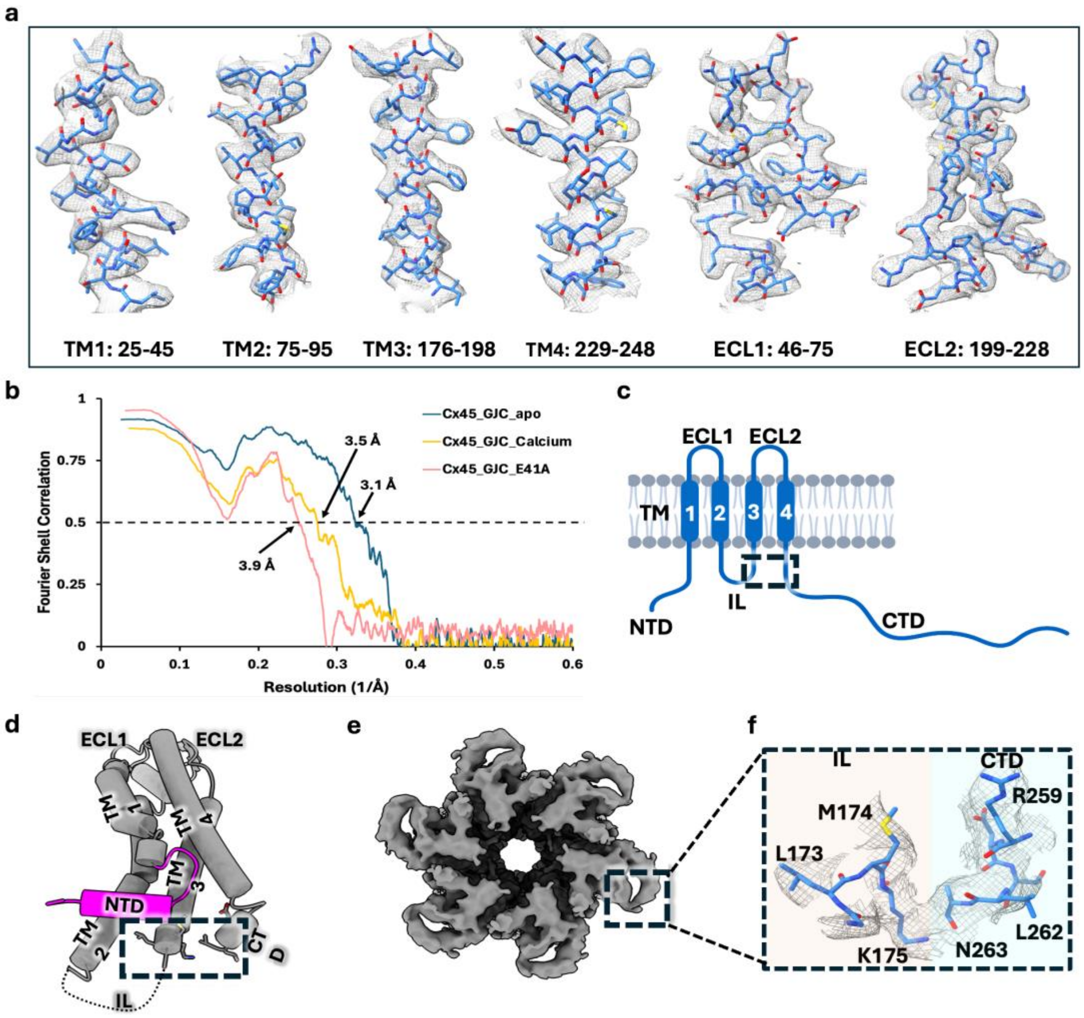


**Supplementary figure 5. Cryo-EM map quality and intracellular loop–C-terminal domain interaction in Cx45. a.** Representative cryo-EM density (grey mesh) and corresponding atomic model (stick representation) illustrating model-to-map agreement in well-resolved regions of the Cx45 gap junction channel. Side-chain density is clearly defined in the transmembrane and extracellular regions, whereas density corresponding to intracellular regions is weaker or absent. **b.** Model–map Fourier shell correlation (FSC) curves for the Cx45 apo (blue), $Ca^{2+}$-bound (wheat), and E41A (salmon) structures. The dashed horizontal line indicates the FSC = 0.5 criterion used to estimate model resolution. **c.** Schematic topology of a Cx45 protomer showing the N-terminal domain (NTD), four transmembrane helices (TM1–TM4), two extracellular loops (ECL1 and ECL2), intracellular loop (IL), and C-terminal domain (CTD). **d.** Tubular representation (tilted view) of a single Cx45 protomer highlighting the domain organization described in panel c. The dashed box indicates the intracellular region highlighted in panels e and f. **e.** Top view of the Cx45 gap junction channel cryo-EM map. The boxed region highlights density corresponding to the intracellular loop (IL) and proximal CTD, suggesting spatial proximity between these regions. **f.** Zoomed-in view of the cryo-EM density (grey mesh) and fitted atomic model showing the interface between the IL and CTD. IL residues L173, M174, and K175 are positioned adjacent to CTD residues R259, L262, and N263. K175 and N263 are located within potential hydrogen-bonding or electrostatic interaction distance, consistent with a possible intramolecular IL–CTD contact.

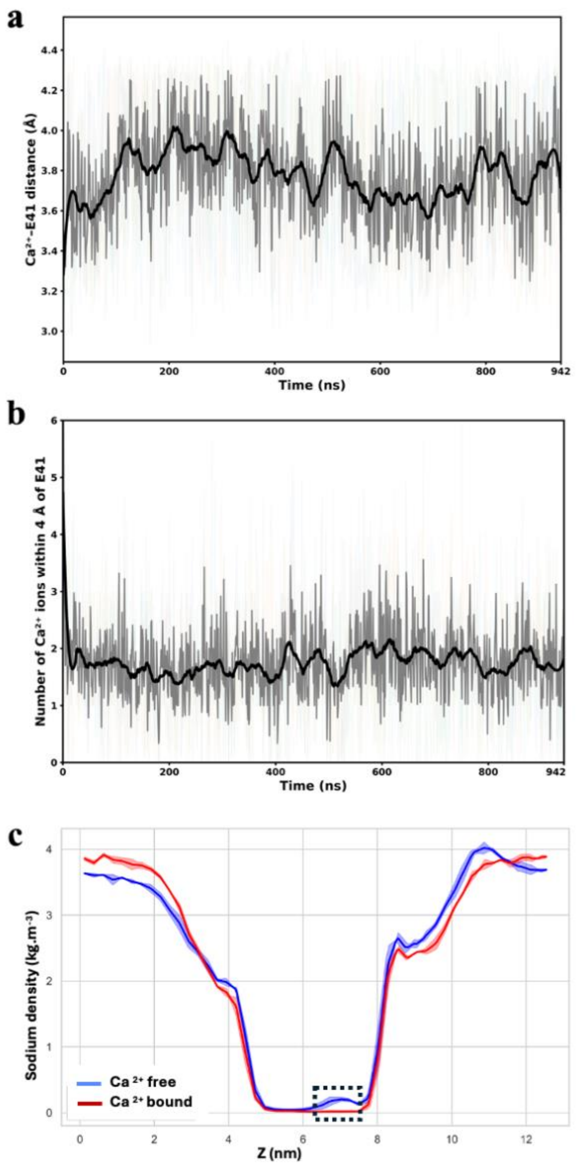


**Supplementary figure 6. Dynamic calcium sampling and sodium exclusion at the E41 neck site in Cx45 simulations. a.** Time evolution of the minimum distance between $Ca^{2+}$ ions and E41 side-chain carboxylate oxygens during molecular dynamics simulations. Calcium frequently approaches the E41 ring within ~3.5–4.0 Å, indicating repeated transient interactions. Individual replicate trajectories are shown as faint lines, with the mean (gray) and a smoothed mean (black) overlaid. **b.** Time evolution of the number of $Ca^{2+}$ ions within 4.0 Å of E41 (per frame). Calcium occupancy fluctuates dynamically around ~1–3 ions, demonstrating recurrent but non-stable binding at the extracellular neck region. **c.** Axial ion density profiles comparing calcium-free (apo, blue) and calcium-bound (red) simulations. Sodium density is elevated under apo conditions but markedly reduced in the calcium-bound state near the extracellular neck region centered on E41 (dashed box), indicating that $Ca^{2+}$ occupancy electrostatically excludes $Na^{+}$ ions from the vestibule.

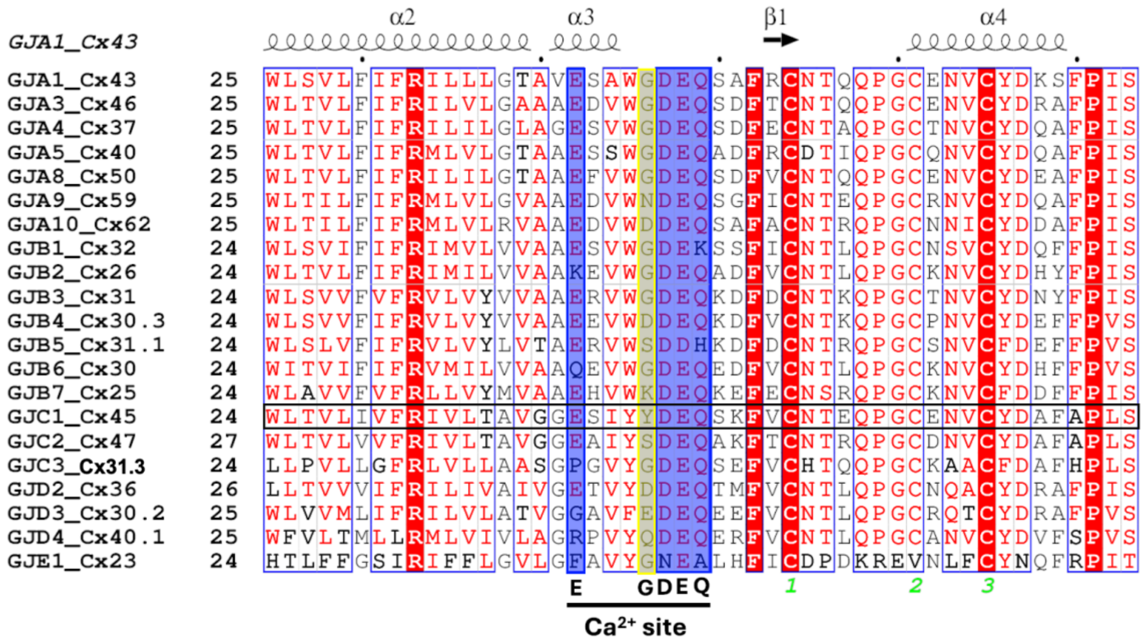

**Supplementary figure 7. Conservation of the extracellular acidic microdomain at the TM1–ECL1 (24-72 residues in Cx45) boundary across human connexins.** Multiple sequence alignment of the TM1–ECL1 region from all 21 human connexins. The conserved acidic motif (E–xxx–GDEQ) is highlighted and marked. In Cx45, E41 corresponds to the first conserved glutamate of this module and is followed by a highly conserved GDEQ sequence. Notably, the glycine residue (highlighted in yellow) preceding the DEQ motif in most connexins is replaced by tyrosine (Y45) in Cx45. The three conserved cysteines (C53, C60, C64 in Cx45) from ECL1 involved in intramolecular disulfide bond formation are marked numerically. (Name discrepancy in literature; GJC3:Cx31.3/Cx30.2/Cx29, GJD3:Cx30.2/Cx31.9).
